# Atomistic Simulations and Network-Based Energetic Profiling of Binding and Allostery in the SARS-CoV-2 Spike Omicron BA.1, BA.1.1, BA.2 and BA.3 Subvariant Complexes with the Host Receptor: Revealing Hidden Functional Roles of the Binding Hotspots in Mediating Epistatic Effects and Long-Range Communication with Allosteric Pockets

**DOI:** 10.1101/2022.09.05.506698

**Authors:** Gennady Verkhivker, Steve Agajanian, Ryan Kassab, Keerthi Krishnan

## Abstract

In this study, we performed all-atom MD simulations of the RBD-ACE2 complexes for BA.1, BA.1.1, BA.2, an BA.3 Omicron subvariants, conducted a systematic mutational scanning of the RBD-ACE2 binding interfaces and analysis of electrostatic effects. The binding free energy computations of the Omicron RBD-ACE2 complexes and comprehensive examination of the electrostatic interactions quantify the driving forces of binding and provide new insights into energetic mechanisms underlying evolutionary differences between Omicron variants. A systematic mutational scanning of the RBD residues determines the protein stability centers and binding energy hotpots in the Omicron RBD-ACE2 complexes. By employing the ensemble-based global network analysis, we proposed a community-based topological model of the Omicron RBD interactions that characterized functional roles of the Omicron mutational sites in mediating non-additive epistatic effects of mutations. Our findings suggested that non-additive contributions to the binding affinity may be mediated by R493, Y498, and Y501 sites and are greater for the Omicron BA.1.1 and BA.2 complexes that display the strongest ACE2 binding affinity among the Omicron subvariants. A network-centric adaptation model of the reversed allosteric communication is unveiled in this study that established a robust connection between allosteric network hotspots and potential allosteric binding pockets. Using this approach, we demonstrated that mediating centers of long-range interactions could anchor the experimentally validated allosteric binding pockets. Through an array of complementary approaches and proposed models, this comprehensive and multi-faceted computational study revealed and quantified multiple functional roles of the key Omicron mutational site R493, R498 and Y501 acting as binding energy hotspots, drivers of electrostatic interactions as well as mediators of epistatic effects and long-range communications with the allosteric pockets.

## 1. Introduction

The enormous body of structural, biochemical, and functional studies established that the mechanism of SARS-CoV-2 infection involves conformational transitions between distinct functional forms of the SARS-CoV-2 viral spike (S) glycoprotein [1–9]. The S protein consists of a conformationally adaptive amino (N)-terminal S1 subunit and structurally rigid carboxyl (C)-terminal S2 subunit, where S1 includes an N-terminal domain (NTD), the receptor-binding domain (RBD), and two structurally conserved subdomains, SD1 and SD2, which coordinate the protein response to binding partners and the host cell receptor ACE2. Conformational plasticity of the SARS-CoV-2 S protein has been extensively examined revealing spontaneous transitions between the closed and open S states that are exemplified by the global movements of the RBDs between the “RBD-down” and “RBD-up” positions [5-15]. Single-molecule Fluorescence (Förster) Resonance Energy Transfer (smFRET) studies characterized the dynamic and kinetics of the SARS-CoV-2 S trimer, showing that the long-range allosteric modulation of the RBD equilibrium can regulate the exposure of the binding site and control ACE2 binding [16-18]. Cryo-EM and X-ray structures of the SARS-CoV-2 S variants of concern (VOC’s) in various functional states and complexes with antibodies unveiled complex and diverse molecular mechanisms that can underlie common and unique dynamics and binding signatures of these proteins [19-28]. These studies demonstrated that the Omicron variants can facilitate the evasion of immune responses induced by vaccination and confer resistance to a wide spectrum of neutralizing antibodies using multiple mutational sites and different mechanisms, whereas preserving and enhancing binding affinity with the ACE2 by modulating interactions in the key energy hotspots. Structural studies of the Omicron variant indicated that evolutionary pressure invokes a complex interplay of thermodynamic factors between mutations that increase affinity for the ACE2 with other RBD modifications that disfavor ACE2 binding but facilitate immune escape [29-33]. These investigations underscored a mechanism in which immune evasion may be a primary driver of Omicron evolution that sacrifices some ACE2 affinity enhancement substitutions to optimize the immune-escaping mutations. The cryo-EM analysis revealed that Omicron mutations can stabilize the Omicron S trimer by strengthening the allosteric interaction network between neighboring protomers and between the S1 and S2 subunits [24]. The crystal structure of the Omicron RBD in complex with human ACE2 determined that the favorable interactions formed by mutated sits S477N, Q493R, Q496S, Q498R, and N501Y to ACE2 could compensate for the loss of polar interactions caused by mutations K417N and E484A [25]. Collectively, these mutations can produce a net enhancing effect on binding of the Omicron RBD to human ACE2 as compared to the original RBD-Wu-Hu-1 strain, suggesting that structural basis for immune evasion while preserving efficient host receptor engagement. The cryo-EM structures of the Omicron variants in different functional states showed that while other variants can promote RBD-up open states and induce immune evasion by mutating common antibody epitopes, the mutant-induced stabilization of the Omicron RBD-down state represents a distinct mechanism that promotes immune evasion by occluding highly immunogenic sites [27]. Consistently, structural studies emphasized a complex and subtle balance of the intermolecular interactions in the Omicron complexes, where substitutions of T478K, Q493R, G496S, and Q498R strengthened the binding of Omicron to ACE2 while mutations K417N and E484A decreased the binding affinity. The cryo-EM study of the full-length S protein of the Omicron variant examined binding and antigenic properties of the Omicron S trimer by bio-layer interferometry showing that the improved binding to ACE2 is mainly due to N501Y, Q493R and Q498R mutations [34]. These structural explorations also indicated that Omicron mutations S477N, T478K and E484A targeting flexible region of the RBM together with K417N may have emerge to ensure the trade-off between a moderate loss of ACE2 binding and the increased neutralization escape potential of the Omicron variant from antibodies. The recent biophysical studies using atomic force microscopy revealed an important role of mechanical stability in mediating immune evasion, showing a synergistic effect of mechanical forces, protein stability effects, and binding interactions that collectively control the virus fitness advantage and immune escape mechanisms [35-37].

By expanding on the initial study of the structures of the Omicron BA.1 RBD in complex with human ACE2 [19], structural basis of the higher binding affinity of ACE2 to currently circulating Omicron subvariants was elucidated in a subsequent investigation which reported the structures of the RBD-ACE2 complexes for BA.1.1, BA.2, and BA.3 variants [38]. This study showed that the Omicron BA.1.1 and BA.2 binding affinities with ACE2 are stronger than the ones for BA.3 and BA.1 subvariants, offering a potential mechanism in which the long-range interaction networks could contribute to the differential binding. The mechanism for enhanced transmission of Omicron BA.2 was examined using structural and biochemical analysis of binding between the human ACE2 and the Omicron BA.2, BA.1, and Wu-Hu-1 variants [39]. This study showed that ACE2 binds to the Omicron BA.2 spike trimer with an affinity which is 11-fold higher than that of the Wu-Hu-1 spike trimer and is nearly 2-fold higher than that with the Omicron BA.1 spike trimer. Structural analysis confirmed that the network of interactions in the BA.2 spike trimer with ACE2 is more extensive and the RBD BA.2 is more stable than the BA.1 [39]. The Omicron subvariants BA.4 and BA.5 featured mutations acquired in the Alpha (N501Y), Beta (K417N, E484K, N501Y), Gamma (K417T, N501Y), and Delta (L452R, T478K), adding E484A and F486V along with T376A, D405N, and R408S modifications that are also present in BA.2. The recent biophysical study described the greater antibody escape from the neutralization of Omicron BA.4 and BA.5 subvariants compared with BA.1 and BA.2, suggesting that unique mutation F486V mutation in BA.4/BA.5 together with L452R and the reversion to the ancestral Q493 are likely to cause more antibody escape [40]. In addition, this study employed surface plasmon resonance (SPR) tools to measure the binding affinity of the Omicron BA.4/5 RBD for ACE2 which was stronger as compared with the ancestral Wu-Hu-1 strain, BA.1, and BA.2 (3-, 3-, and 2-fold, respectively) likely due to an increase in binding half-life and the improved electrostatic complementarity. A systematic antigenic analysis of the Omicron subvariants showed that L452R found in both BA.2.12.1 and BA.4/5 can facilitate escape from classes II/III antibodies, while F486V mutation from BA.4/5 promotes escape from classes I/II antibodies and compromises the ACE2 affinity [41]. This study suggested that F486V in BA.4 and BA.5 allows for antibody evasion while R493Q reversion allows to regain binding fitness and may eve lead for BA.4/BA.5 to have a slightly higher affinity for ACE2 as compared to other Omicron subvariants.

A tour-de-force structural and biochemical study reported cryo-EM structures of the S trimers for BA.1, BA.2, BA.3, BA.2.12.1, BA.2.13 and BA.4/BA.5 subvariants of Omicron [42]. The binding affinities of the Omicron variants with the ACE2 receptor determined by SPR showed a decreased binding affinity for the BA.4/BA.5 subvariants, also revealing that the Omicron BA.2 displayed slightly higher binding affinities than the other Omicron variants. Using biophysical analysis, this study suggested that the F486V mutation in BA.4/BA.5 is a primary contributor for a moderately decreased ACE2 binding affinity, while the reversion R493Q can only marginally impact binding interactions [42]. In-depth functional analysis of 27 potent RBD-binding antibodies isolated from vaccinated volunteers following breakthrough Omicron-BA.1 infection revealed neutralization differences between sublineages, particularly showing that BA.2 has a slightly higher ACE2 affinity than BA.1 and slightly reduced neutralization [43]. Structure-functional studies of the Omicron BA.1, BA.2, BA.2.12.1, BA.4 and BA.5 subvariants showed the increased ACE2 binding affinity, decreased fusogenicity, and stronger evasion of neutralizing antibody responses as compared to the Wu-Hu-1 and Delta strains, confirming that the compounded effect of the enhanced ACE2 receptor binding and stronger immune evasion may have contributed to the rapid spread of these Omicron sublineages [44]. Using bio-layer interferometry and SPR techniques, this investigation quantified the greater binding affinity of the BA.2 as compared to the original Wu-Hu-1 strain and original Omicron BA.1 variant, but in some contrast to other studies reported the greatest binding affinity of the BA.4/BA.5 RBD [44]. Hence, multiple structural studies revealed generally similar and high binding affinities across all the Omicron subvariants, while showing marginally better binding profile for the BA.1.1 and especially BA.2 subvariants.

Computer simulations provided important atomistic and mechanistic advances into understanding the dynamics and function of the SARS-CoV-2 S proteins [45-57]. All-atom MD simulations of the full-length SARS-CoV-2 S glycoprotein embedded in the viral membrane, with a complete glycosylation profile, were performed by Amaro and coworkers, providing an unprecedented level of detail about the conformational landscapes of the S proteins in the physiological environment [45]. Another landmark study from this laboratory reported 130 µs of weighted ensemble simulations of the fully glycosylated S ectodomain and statistical characterization of more than 300 kinetically unbiased RBD-opening pathways [47]. Using distributed cloud-based computing, large scale MD simulations of the viral proteome observed dramatic opening of the S protein complex, predicting the existence of several cryptic epitopes in the S protein [50]. MD simulations of the S-protein in solution and targeted simulations of conformational changes between the open and closed forms revealed the key electrostatic interdomain interactions mediating the protein stability and kinetics of the functional spike states [56]. Using the replica-exchange MD simulations with solute tempering of selected surface charged residues, the conformational landscapes of the full-length S protein trimers were investigated, unveiling previously unknown cryptic pockets and the meta-stable intermediate states [57]. A number of computational studies employed atomistic simulations and binding energy analysis to examine the interactions between the S-RBD Omicron and the ACE2 receptor [58-62]. All-atom MD simulations of the S Omicron trimer and the Omicron RBD–ACE2 complexes suggested that the Omicron mutations may have evolved to inflict a greater infectivity using a combination of more efficient RBD opening, the increased binding affinity with ACE2, and optimized capacity for antibody escape [58].

All-atom MD simulations of the BA.1 and BA.2 RBD complexes with ACE2 in the presence of full-length glycans revealed a more dispersed interaction network and the increased number of salt bridges and hydrophobic interactions with ACE2 compared to the original Wu-Hu-1 RBD [59]. Computational studies demonstrated that Omicron S RBD binding to the ACE2 receptor can be primarily determine by long-range electrostatic forces also suggesting the evolution of the electrostatic surface between VOCs by gradual accumulation of positive surface charges in the RBD [61]. By providing a quantitative assessment of the antibody escape, this study showed that Omicron variant may promote resistance to the most neutralizing antibodies by modulating the electrostatic interactions. These observations were confirmed in another computational study in which atomistic MD simulations examined structural and dynamic effects of the Omicron RBD-ACE2 complex using fully glycosylated models of the mutant variants and demonstrated the improved electrostatic contribution to binding as well as a considerable stabilization of the binding interface [62]. MD simulations of the Omicron RBD binding with ACE2 suggested that strong electrostatic interactions can promote the increased infectivity and transmissibility compared to other strains, also showing that K417N, G446S, and Y505H mutations can decrease the ACE2 binding, while S447N, Q493R, G496S, Q498R, and N501Y mutations improve binding affinity with the host receptor [63]. Sequence and structure-based computational analysis of the Omicron BA.2 showed the reversed G446/G496 combination makes BA.2 more stable than Omicron BA.1 featuring S446/S496 mutations. The study concluded that Omicron BA.2 subvariant may have evolved to maintain ACE2 binding similar to the original strain while promoting more efficient antibody evasion highlighting how complex pattern of mutations balances these conflicting factors [64]. By employing the conformational ensembles of the S-RBD Omicron variant complexes with ACE2 we recently performed simulations and mutational scanning of the interfacial RBD residues showing that N501Y is the critical binding affinity hotspot in the S Omicron RBD complex with ACE2, while hotspots Q493R, G496S, Q498R anchor the key interfacial clusters responsible for binding with ACE2 [65]. By assessing over 700 mutant complexes, the recent study revealed that high-affinity RBD mutations (including N440K, S443A, G476S, E484R, G502P) tend to cluster near known human ACE2 recognition sites supporting the view that combinatorial mutations in SARS-CoV-2 can develop in sites amenable for non-additive enhancements in binding and antibody evasion. simultaneously maintain high-affinity binding to ACE2 and evade antibodies (e.g., by N440K, L452R, E484K/Q/R, K417N/T) [66]. The effect of nonadditive, epistatic relationships among RBD mutations was assessed using protein structure modeling by comparing the effects of all single mutants at the RBD-ACE2 interfaces for the Omicron variants, showing that structural constraints on the RBD can curtail the virus evolution for a more complete vaccine and antibody escape [67]. A systematic analysis of the epistatic effects in the S-RBD proteins was undertaken in an illuminating deep mutational scanning study measuring the impacts of all amino acid mutations in the RBD in the Wu-Hu-1, Alpha, Beta, Delta, and Eta variants [68]. This study showed that N501Y causes significant epistatic shifts in the effects of mutations at Q498 as well as RBD residues 446-449 and 491-496 which take place in the absence of any appreciable structural changes between the RBD variants. The evolutionary and functional studies suggested that the emergence of the Omicron variants can be determined by multiple fitness trade-offs including the immune escape, binding affinity for ACE2, conformational plasticity, protein stability and allosteric modulation [69-71]. Our previous studies revealed that the SARS-CoV-2 S protein can function as an allosteric regulatory machinery that can exploit the intrinsic plasticity of functional regions controlled by stable allosteric hotspots to modulate specific regulatory and binding functions [72-78]. However, the dynamic and energetic details quantifying the balance and contribution of these factors, particularly the role of long-range interactions in binding of the Omicron subvariants with ACE2 and antibodies remain mechanistic and scarcely characterized.

In this study, we perform all-atom MD simulations of the RBD-ACE2 complexes for BA.1 BA.1.1, BA.2, an BA.3 Omicron subvariants, and conduct a systematic mutational scanning of the RBD-ACE2 binding interfaces and assessment of the electrostatic effects using equilibrium ensembles and robust computations of the binding free energy changes. Through a detailed analysis of the dynamics and intermolecular interactions, we characterize the fundamental commonalities and differences in the interaction profiles for the Omicron RBD subvariants. The rigorous binding free energy computations of the Omicron RBD-ACE2 complexes and comprehensive examination of the electrostatic interactions quantify the driving forces of binding and provide new insights into energetic-driven evolutionary differences between Omicron variants. In addition, we perform a systematic mutational scanning of the RBD residues that determines the protein stability centers and binding energy hotpots in the Omicron RBD-ACE2 complexes. By employing the ensemble-based global network analysis, we propose a community-based topological model of the inter-residue connectivity in the Omicron RBD complexes to examine the role of Omicron mutational sites in mediating potential non-additive epistatic effects of mutations. Our findings suggest that the extent of non-additive contributions to the binding affinity may be greater for the Omicron BA.1.1 and especially BA.2 complex that also featured the strongest binding affinity among the Omicron subvariants. We also propose a new network-centric adaptation of the reversed allosteric communication approach to identify allosteric mediating hotspots and infer this analysis to characterize the distribution of allosteric binding pockets on the RBD. We show that this approach can successfully identify the experimentally known allosteric binding sites that are anchored by the predicted allosteric hotspots. The results showed that using reversed allosteric mapping, we can identify the experimentally validated conserved allosteric binding pocket and characterize allosteric cross-talk between allosteric binding site and RBD-ACE2 binding interface. Through a battery of complementary modeling approaches, this comprehensive and multi-faceted computational study revealed and quantified multiple functional roles of the key Omicron mutational site R493, R498 and Y501 acting as binding energy hotspots, drivers of electrostatic interactions as well as mediators of epistatic effects and long-range communications with the allosteric pockets.

## 2. Results and Discussion

### 2.1. Atomistic MD Simulations Reveal Common and Distinct Signatures of Conformational Dynamics and Interaction Patterns in the ACE2 Complexes with the Omicron RBD Variants

We performed a detailed structural and dynamic analysis of the Omicron RBD BA.1, BA.1.1, BA.2 and BA.3 complexes with the host receptor human ACE2 (hACE2) (Figure 1). A total of 12 mutations (G339D, S373P, S375F, K417N, N440K, S477N, T478K, E484A, Q493R, Q498R, N501Y, and Y505H) are shared among the BA.1 and BA.2 variants. In the RBD, BA.1 contains unique mutations S371L, G446S, and G496S while BA.1.1 subvariant features a unique R346K mutation, BA.2 carries S371F, T376A, D405N, and R408S mutations, and BA.3 has S371F and D405N mutations (Table 1). Structural analysis of the RBD binding epitopes in the complexes revealed a very similar composition of the interacting residues for all studied Omicron subvariants and virtually identical topography of the binding interface (Figure 1). In the Omicron RBD BA.1-hACE2 complex, the signature mutational sites G446S, T478K, E484A, Q493R, G496S, Q498R, N501Y, and Y505H belong to the binding epitope but K417N site is no longer within the interacting distance from the host receptor (Figure 1A,B).

**Figure 1.**
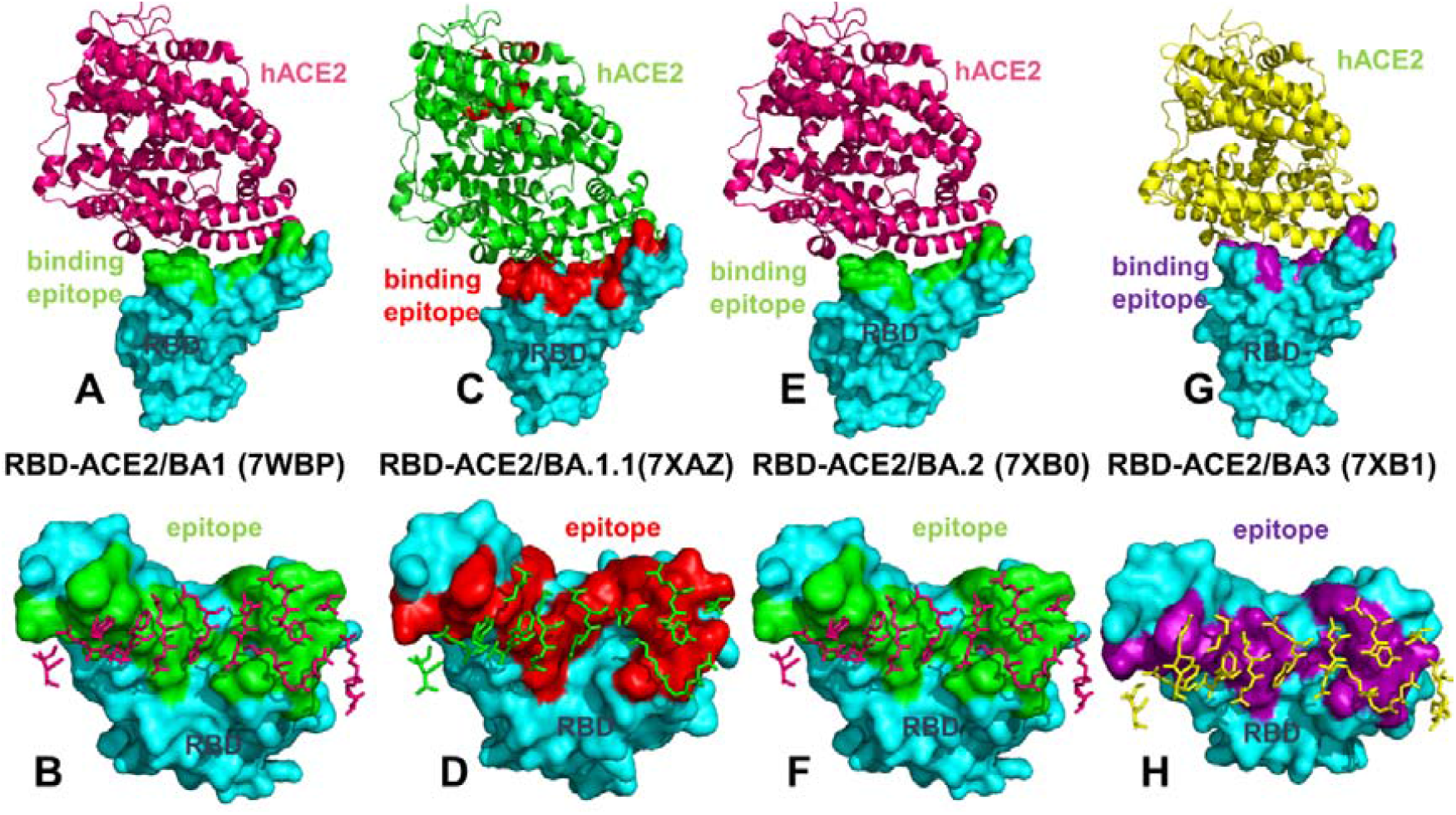
Structural organization and binding epitopes of the SARS-CoV-2-RBD Omicron BA.1, BA.1.1, BA.2 and BA.3 complexes with human ACE enzyme. (A) The cryo-EM structure of the Omicron RBD BA.1-ACE2 complex (pdb id 7WBP). The RBD is shown in cyan-colored surface and the bound ACE2 enzyme is in pink ribbons. (B) The RBD-BA.1 binding epitope is highlighted in green surface. The ACE2 binding residues are shown in pink sticks. (C) The cryo-EM structure of the Omicron RBD BA.1.1-ACE2 complex (pdb id 7XAZ). The RBD is shown in cyan-colored surface and the bound ACE2 enzyme is in green ribbons. (D) The RBD-BA.1.1 binding epitope is shown in red surface. The ACE2 binding residues are shown in green sticks. (E) The cryo-EM structure of the Omicron RBD BA.2-ACE2 complex (pdb id 7XB0). The RBD is shown in cyan-colored surface and the bound ACE2 enzyme is in pink ribbons. (F) The RBD-BA.2 binding epitope is shown in green surface and the ACE2 binding residues are shown in pink-colored sticks. (G) The cryo-EM structure of the Omicron RBD BA.3-ACE2 complex (pdb id 7XB1). The RBD is shown in cyan-colored surface and the bound ACE2 enzyme is in green ribbons. (H) The RBD-BA.3 binding epitope is shown in purple-colored surface and the ACE2 binding residues are shown in green sticks.

**Table 1.** Mutational landscape of the Omicron subvariants in the S-RBD.

| SARS-CoV-2 Omicron Variant | Mutational landscape of the RBD |
| --- | --- |
| BA.1 | G339D, S371L, S373P, S375F, K417N, N440K, G446S, S477N, T478K, E484A, Q493R, G496S, Q498R, N501Y, Y505H |
| BA.1.1 | G339D, **R346K**, S371L, S373P, S375F, K417N, N440K, G446S, S477N, T478K, E484A, Q493R, G496S, Q498R, N501Y, Y505H |
| BA.2 | G339D, **S371F**, S373P, S375F, **T376A**, **D405N**, **R408S**, K417N, N440K, S477N, T478K, E484A, Q493R, Q498R, N501Y, Y505H |
| BA.3 | G339D, **S371F**, S373P, S375F, **D405N**, K417N, N440K, G446S, S477N, T478K, E484A, Q493R, Q498R, N501Y, Y505H |
| BA.4/BA.5 | G339D, **S371F**, S373P, S375F, **T376A**, **D405N**, **R408S**, K417N, N440K, **L452R**, S477N, T478K, E484A, **F486V**, Q493R, Q498R, N501Y, Y505H |

Structural analysis of the Omicron BA.1 complex unveiled a salt-bridge between R498 and K353 of hACE2 as well as a hydrogen bond between S496 and K353. The binding epitope for the Omicron BA.1.1 complex is similar to that of BA.1 but additionally featured K417N and V445 residues (Figure 1C,D). The binding epitope in the BA.1.1 subvariant highlighted the apparent strengthening in the middle part of the binding interface, owing to a stronger interaction network mediated by R493 with K31, E35 and D38 of hACE2. The two mutations in BA.2 G446 and G496 reverted to the Wu-Hu-1 residues. Despite minor conformational changes in the loops harboring G446/496 sites resulting from presence of R498 and Y501 mutations in the BA.2 subvariant, the binding epitope topography remain largely intact (Figure 1E,F). A modest enlargement at the edges of the binding epitope in the RBD BA.2 complex can be noticed due to additions of R403, N417, V445, and Q506 residues. BA.3 includes D405N but not G496S mutations. Neither of these residues is involved in the intermolecular contacts with hACE2. However, the topography of the binding epitope in the RBD BA.3-hACE2 complex highlighted a considerable similarity to other Omicron subvariants. This preliminary analysis indicated that structural differences between the Omicron RBD complexes are extremely small and cannot alone explain subtle differences in the binding affinity with the host receptor. The structural and functional similarities between Omicron subvariants were also apparent in the antibody binding profiles, showing that BA.1.1, BA.2, and BA.3 showed immune evasion spectra as BA.1 [79].

By assuming that the functional and binding differences between Omicron variants may be determined by dynamic signatures and variations in stability of the intermolecular interactions, we performed ten independent all-atom MD simulations (500 ns for each simulation) of the studied protein systems (Table 2). Conformational dynamics profiles of the RBD and hACE2 residues are described using the root mean square fluctuations (RMSF) obtained from MD simulations (Figure 2). The RBD has two subdomains, where the flexible RBM with a concave surface is involved in direct interaction contacts with hACE2 (Figure 1). The second subdomain is a structured five-stranded antiparallel β-sheet core region that functions as a stable core scaffold for the RBM. The conformational mobility distributions for the Omicron RBD complexes displayed several deep local minima corresponding to residues 374-377 and the RBD core residue cluster (residues 396-403) as well as interfacial RBD positions that are involved in the contacts with the hACE2 receptor (residues 445-456 and 496-505 of the binding interface) (Figure 2A). The observed structural stability of the core RBD regions was also seen in our earlier simulation studies of the original RBD Wu-Hu-1 and Omicron complexes [65], indicating that these segments remain relatively rigid in all Omicron variant complexes with hACE2. Noteworthy, the most stable RBD positions included important hydrophobic stability centers F400, I402, F490, Y453, L455, A475, and Y489 (Figure 2A). Some of these hydrophobic RBD positions (Y453, L455, A475 and Y489) are also involved in the favorable interfacial contacts with hACE2 and constitute an indispensable conserved region of the RBD-hACE2 interface. The RBD binding loop containing residues 474–486 experienced moderate fluctuations in Omicron RBD-hACE2 complexes, while another distal allosteric loop (residues 358–376) showed an appreciable mobility with the exception of S371L/F, S373P and S375F that displayed reduced fluctuations in the complexes (Figure 2A).

**Table 2.** Structures of the Omicron RBD-hACE2 complexes examined in this study.

| PDB | System | Per simulation | # simulations |
| --- | --- | --- | --- |
| 7WBP | Omicron RBD BA.1-hACE2 | 500 ns | 10 |
| 7XAZ | Omicron RBD BA.1.1-hACE2 | 500 ns | 10 |
| 7XB0 | Omicron RBD BA.2-hACE2 | 500 ns | 10 |
| 7XB1 | Omicron RBD BA.3-hACE2 | 500 ns | 10 |

**Figure 2.**
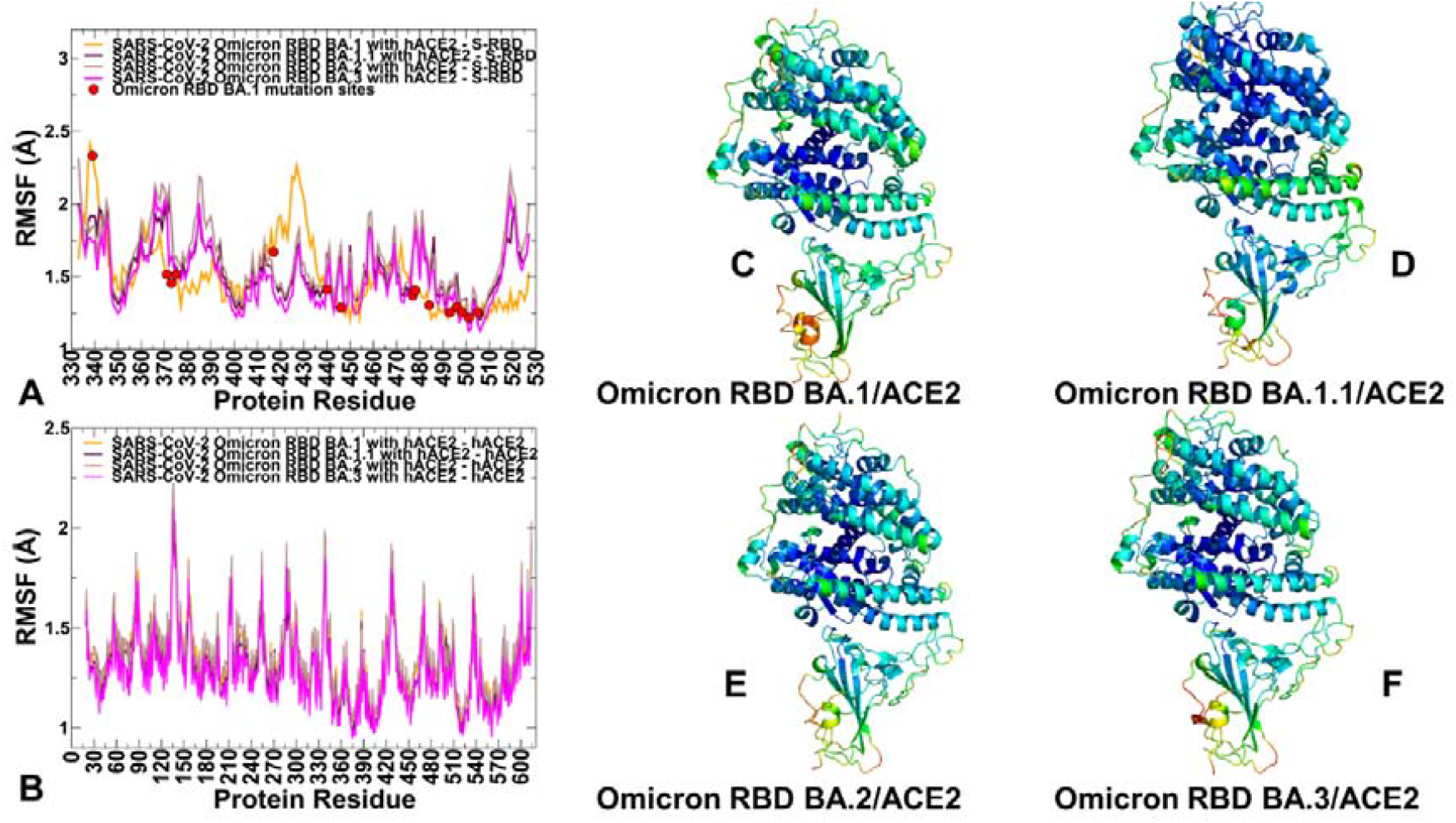
Conformational dynamics profiles obtained by averaging results from 10 independent MD simulations of the Omicron RBD BA.1, BA.1.1, BA.2 and BA.3 complexes with hACE2. (A) The RMSF profiles for the RBD residues obtained from MD simulations of the RBD BA.1-hACE2 complex, pdb id 7WBP (in green lines), RBD BA.1.1-hACE2 complex, pdb id 7XAZ (in maroon lines), RBD BA.2-hACE2 complex, pdb id 7XB0 (in light brown lines), and RBD BA.3-hACE2 complex, pdb id 7XB1 (in magenta lines), The positions of Omicron mutational sites are highlighted in red-colored filled circles. (B) The RMSF profiles for the hACE2 residues obtained from MD simulations of the Omicron BA.1, BA.1.1, BA.2 and BA3 complexes. The ACE2 dynamics profiles are shown respectively in green, maroon, light brown and magenta lines for BA.1, BA.1.1, BA.2 and BA3 complexes respectively. Structural maps of the conformational profiles are obtained from MD simulations of Omicron RBD variant complexes. Conformational mobility maps for the Omicron RBD BA.1-hACE2 complex (C), the Omicron RBD BA.1.1-hACE2 complex (D), the Omicron RBD BA.2-hACE2 complex (E), and the Omicron RBD BA.3-hACE2 complex (F). The structures are shown in ribbons with the rigidity-flexibility sliding scale colored from blue (most rigid) to red (most flexible). The positions of sites targeted by Omicron mutations are shown in spheres colored according to their mobility level.

Mutation S373P is identical and relatively conservative, S371L mutation is shared between BA. 1 and BA.1.1 while S371F mutation is common to BA.2, BA.3, BA.4 and BA.5 subvariants. The emergence of Omicron mutations in the stable scaffolding subdomain of the RBD indicated that these variations may affect the mechanical stability of the RBD. MD simulations showed a considerable stabilization in these positions, particularly for the BA.1 and BA.1.1 complexes (Figure 2A). Despite generally common dynamic profiles of the RBD in all Omicron-hACE2 complexes, we noticed that the RBD-BA.2 and RBD-BA.3 displayed smaller fluctuations in the residues 400-440, while this RBD segment experienced more significant thermal displacements in the RBD-BA.1 complex (Figure 2A). The portion of the RBM region (residues 446–494) that provides the contact interface with ACE2 also displayed appreciably smaller movements and becomes largely stabilized in the Omicron RBD complexes (Figure 2A). The mobile flexible RBM loops (residues 473-487) appeared to be partly constrained in the RBD-BA.1 complex but remained flexible in the other Omicron RBD complexes (Figure 2A). Noticeably, residues 470-480 retain a certain degree of flexibility despite contributing appreciably to the intermolecular interface. At the same time, the reduced RMSF values were noticeable for the RBD residues 480-505 that are involved in the key interactions with hACE2 and become largely stabilized for all complexes. In particular, the RMSF profile highlighted a considerable stabilization of the binding interface residues Q493R, Q498R, N501Y, Y505H that are shared between Omicron BA.1, BA.1.1, BA.2, BA.3, BA.4 and BA.5 (Figure 2A) suggesting that this region of the binding interface can contribute decisively to the stability and binding affinity of the Omicron RBD-hACE2 complexes.

By mapping the positions of the Omicron mutations onto the profile, one could immediately notice that most of the mutational sites displayed only moderate or small fluctuations with the exception of G339D position that is highly flexible in the RBD-BA.1 complex but becomes mobile in other complexes (Figure 2A). Another interesting observation of the comparative dynamic analysis is the conformational mobility of K417N in the RBD-BA.1 complex and the increased flexibility of S477N and T478K Omicron sites in the BA.1.1, BA.2 and BA.3 complexes. The conformational dynamics profile of the hACE2 receptor showed a similar and strong stabilization of the interfacial helices on ACE2, indicating that dynamics signatures of the bound hACE2 receptor remain largely conserved across all Omicron RBD complexes (Figure 2B). Structural mapping of the conformational dynamics profiles further highlighted similarities between complexes, while showing a subtle but visible increase in the RBD stability of the BA.1.1 and BA.2 subvariants (Figure 2C-F). According to Our analysis, the largest increase in stability of the RBD-hACE2 complexes may be induced by Omicron BA.1.1 mutations. The results of simulations also indicated that both structural and dynamic signatures of the Omicron RBD subvariants remain largely conserved and subtle differences in their binding affinity with hACE2 may result from subtle rearrangements and cumulative effects of many intermolecular contacts rather than being attributed to the mutational changes of several specific RBD residues.

### 2.2. Ensemble-Based Analysis of the Intermolecular Interactions in the Omicron RBD-hACE2 Complexes : Quantifying Common and Unique Energetic Signatures of the Omicron RBD Variants

Using conformational ensembles of the Omicron RBD-hACE2 complexes, we performed a detailed statistical analysis of the intermolecular contacts that revealed several fundamental commonalities and differences in the interaction profiles for the Omicron RBD subvariants (Figures 3,4). The RBD-hACE2 contacts with the occupancy > 70% in the MD trajectories were considered as long-lived stable interactions and were recorded for this analysis. In particular, we examined the ensemble-averaged number of distinct ACE2 residues making stable contacts with each of the RBD binding residues. Using this parameter, the density and strength of the pair-wise RBD-hACE2 interactions can be determined and systematically compared between Omicron RBD subvariants. The RBD residues are generally organized in several groups based on their conservation and respective roles in binding with the host receptor. One of these groups includes identical residues such as Y453, N487, Y489, T500, and G502 and homologous positions (Y449/F/H, F456/L, Y473/F, F486/L, and Y505H), while other group includes more diverse residues undergoing various modifications such as G446/S/T, L455/S/Y, A475/P/S, G476/D, G496/S, K417/V/N/R/T, E484/K/P/Q/V/A, Q493/N/E/R/Y, Q498/Y/H/R, and N501/Y/T/D/S [28]. Our analysis confirmed that the conserved residues from the first group make consistent and similar interactions in all Omicron variants, indicating that these RBD positions can act as molecular determinants of the RBD stability and binding affinity (Figures 3,4). The modifications of residues from the other group such as G446S, G496S, K417N, E484A, 493R, Q498R, and N501Y are utilized in the Omicron variants and could modulate binding affinity with the hACE2.

**Figure 3.**
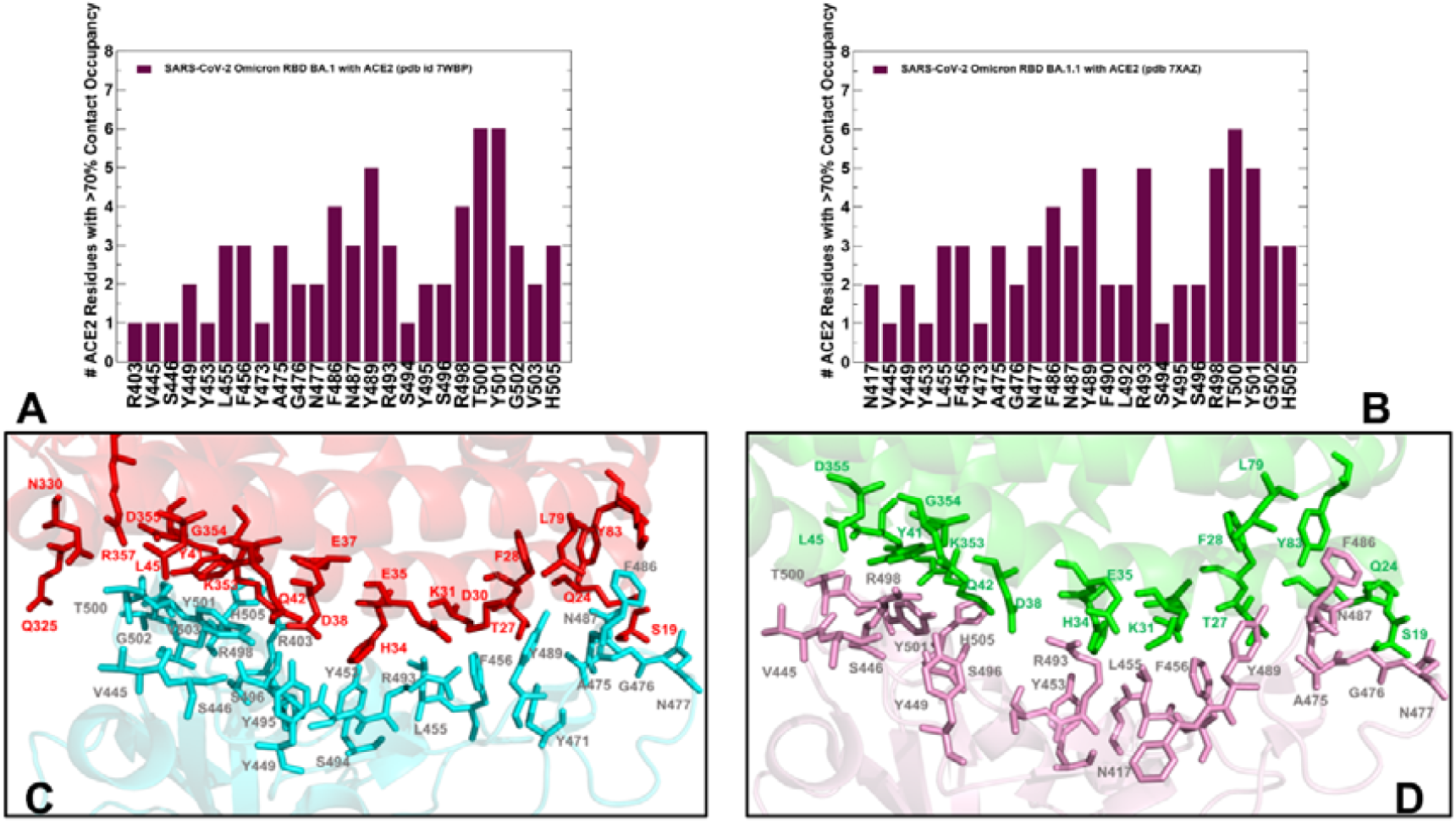
The distributions of the intermolecular contacts obtained by averaging results from 10 independent MD simulations of the Omicron RBD BA.1 and BA.1.1 complexes with hACE2. The ensemble-average number of distinct hACE2 residues making stable intermolecular contacts (> 70% occupancy) with the RBD residues obtained from MD simulations of the RBD BA.1-hACE2 complex, pdb id 7WBP (A) and the RBD BA.1.1-hACE2 complex, pdb id 7XAZ (B). Structural mapping of the intermolecular RBD BA.1-hACE2 interface. The RBD binding residues are shown in cyan sticks and the hACE2 binding residues are in red sticks (C). Structural mapping of the intermolecular RBD BA.1.1-hACE2 interface. The RBD binding residues are shown in cyan sticks and the hACE2 binding residues are in green sticks. The intermolecular binding interface residues are annotated.

**Figure 4.**
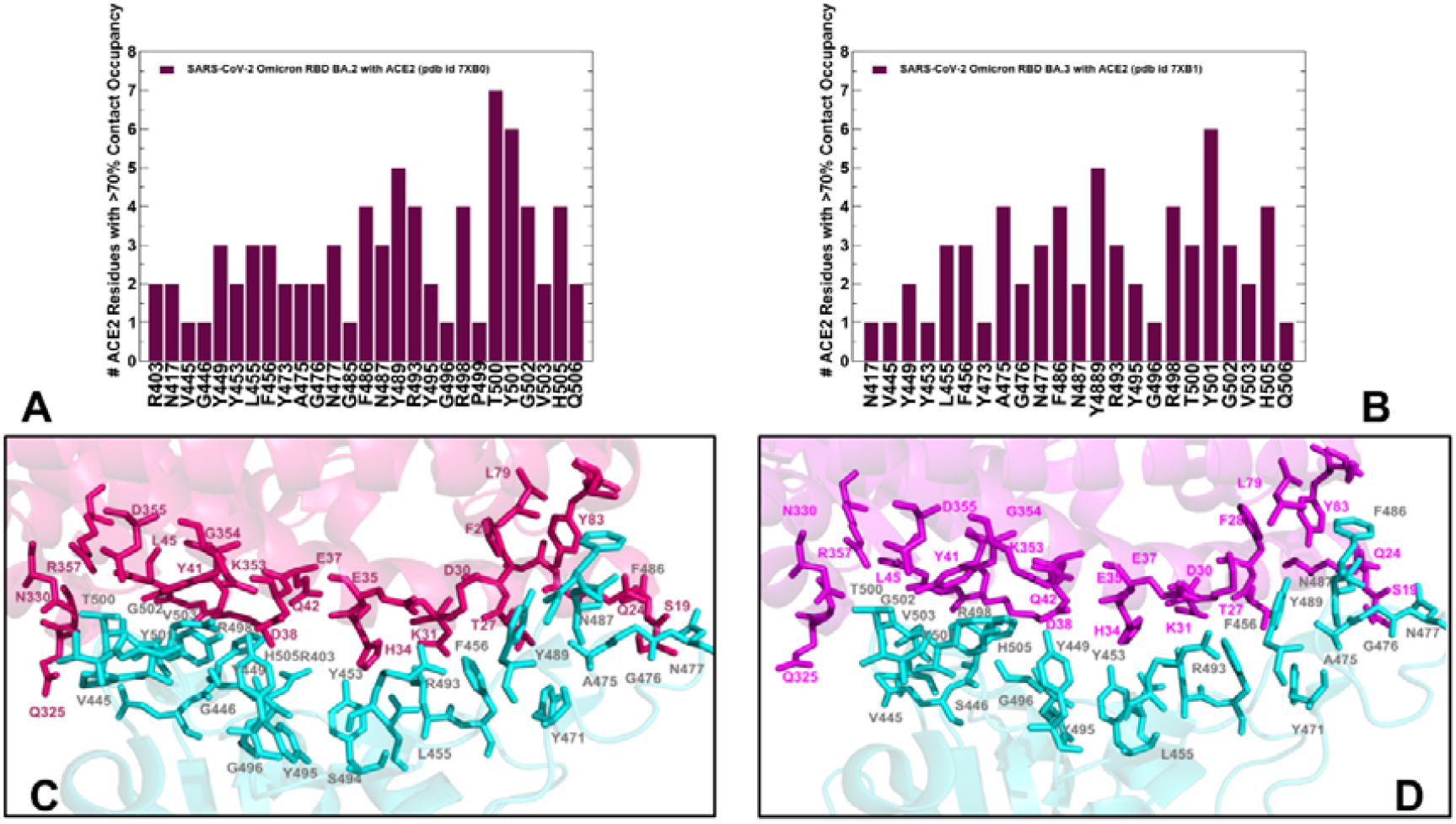
The distributions of the intermolecular contacts obtained by averaging results from 10 independent MD simulations of the Omicron RBD BA.2 and BA.3 complexes with hACE2. The ensemble-average number of distinct hACE2 residues making stable intermolecular contacts (> 70% occupancy) with the RBD residues obtained from MD simulations of the RBD BA.2-hACE2 complex, pdb id 7XB0 (A) and the RBD BA.3-hACE2 complex, pdb id 7XB1 (B). (C) Structural mapping of the intermolecular RBD BA.1-hACE2 interface. The RBD binding residues are shown in cyan sticks and the hACE2 binding residues are in dark, pink-colored sticks. (D) Structural mapping of the intermolecular RBD BA.1.1-hACE2 interface. The RBD binding residues are shown in cyan sticks and the hACE2 binding residues are in magenta-colored sticks. The intermolecular binding interface residues are annotated.

We first compared the distributions for the RBD BA.1 and BA.1.1 complexes (Figure 3). The analysis reinforced the notion that the interaction contacts formed by conserved residues N487, Y489, T500 contribute significantly to the binding (Figure 3A,B). The stable contacts along the binding interface between the RBD and hACE2 are distributed over two patches [19,38].

The patch 1 in all complexes contains hACE2 residues S19, Q24, H34, E35, and Y83 which interact with the RBD residues. In the patch 1 of the RBD BA.1 and BA.1.1 complexes, Q24 of hACE2 interacts with N487 of RBD, Y83 of hACE2 forms hydrogen bonds with Y489 and N487 of RBD, and A475 from RBD makes stable contacts with S19, Q24 and T27 (Figure 3, Table S1). In addition, a segment of the patch 1 in the middle of the binding interface contains H34 from hACE2 contacting Y453 of RBD, E35 of hACE2 forming a salt bridge with R493 from RBD, F456 making stable contacts with T27, E30, K31 and F486 from RBD packing against F28, L79, M82, and Y83 of hACE2 (Figure 3, Table S1). Despite similar distribution profiles of the interfacial contacts and interacting residues in the RBD BA.1 and BA.1.1 complexes, our analysis pointed to several important differences. The major distinctions of the RBD BA.1.1 interactions are : (a) a noticeable broadening of the RBD binding interface and an addition of the N417, F490, L492 sites; and (b) the increased number of stable contacts and interacting hACE2 positions for R493 position, including a hydrogen bond with D38 and a salt bridge with S496 (Figure 3, Table S1). The interfacial contacts of R493 with K31, H34, E35, and D38 in the RBD BA.1.1 complex form a very stable electrostatic intermolecular network that appeared to be unique for this Omicron subvariant (Figure 3). In addition, new stable interfacial contacts with hACE2 are formed by F490 and L492 expanding the spectrum of RBD residues in the patch 1. The patch 2 of the binding interface includes D38, Y41, Q42, L45, G352, K353, G354, and D355 of hACE2 and forms the extensive network of interactions and hydrogen bonds with the RBD residues G496S, R498, T500, Y501, G502, V503 and H505 (Figure 3, Table S1). In particular, R498 interacts with D38, Y41, Q42 and L45 ACE2 residues, Y501 maintains stable contacts of high occupancy with D38, Y41, G352, K353, G354, and D355 while H505 makes stable contacts with E37, K353, and G354 residues. In all Omicron RBD complexes, Y501 and H505 together with R498 form a stable cluster that restricts movements of the RBD residues in this region and can stabilize the binding interface. These stable interactions result in the increased rigidification of the negatively charged ACE2 residues D30, E35, E37, and D38. By averaging time-dependent evolution of the interaction contacts over 10 independent 500ns MD simulations, we monitored the occupancy of important salt bridges and interfacial hydrogen bonds (Table S2). The results showed a markedly increased stability of salt bridges in the Omicron RBD BA.1.1 complex. Indeed, the occupancies of major salt bridges R403-E37 (65%/78%), K440-E329 (31%/56%), R493-E35 (77%/88%), R493-D38(26%/95%) and R498-D38 (59%/97%) revealed the increased stability of these interactions in the BA.1.1 complex (Table S2).

We observed that the patch 2 of the RBD (G496S, R498, T500, Y501, G502, V503 and H505) is enriched in the RBD BA.2-hACE2 complex, forming the largest number of diverse contacts and establishing the strongest interaction network (Figure 4A,C). The distributions highlighted an extensive network of the ACE2 residues interacting with T500 and Y501 positions (Figure 4A). In the RBD BA.2 complex, T500 of RBD forms long-lived stable interactions with Y41, L45, G326, Q330, K353, G354, D355 and R357 while a critical for binding Y501 residue is engaged in the favorable stable contacts with D38, Y41, K353, G354 and D355 (Figure 4A,C). Common to all Omicron RBD-hACE2 complexes are strong π-π interactions formed with Y41 and an additional π-cation interaction with K353 (Tables S1, S2). Although these important contacts are stable in all complexes, the occupancy of these specific interactions is marginally higher in the BA.1.1 and BA.2 complexes. The results agree with related studies which combined MD simulations and single-molecule force microscopy (SMFS) to show that the N501Y mutation-induced π-π and π-cation interactions could explain the changes observed by force microscopy [80].

Structural mapping of the interfacial residues involved in the high occupancy interactions in the BA.2 complex illustrates a dense and more extended network in the patch 2 (Figure 4C) as compared to the RBD BA.1 (Figure 3C) and BA.1.1 complexes (Figure 3D). Our analysis showed that the RBD-BA.2 engages in the patch 2 the largest number of ACE2 residues (Q325, G336. G352, K353, G354, D355, an R357) (Figure 4C, Table S1). Our analysis showed that among the BA.1 RBD/hACE2, BA.2 RBD/hACE2, and BA.3 RBD/hACE2 complexes, the BA.2 RBD/hACE2 binding interface has the largest number of highly stable intermolecular contacts and hydrogen bonds (Figure 4, Tables S1,S2). These findings are consistent with the higher binding affinity of the RBD BA.2 as compared with BA.1 and BA.3 [38]. Indeed, the salt bridges, hydrogen bonds and hydrophobic interactions were very stable in the RBD BA.2 complex and displayed the higher occupancy of the favorable contacts as compared to other Omicron RBD-hACE2 complexes (Table S2).

We found that hydrogen bonds T500-D355 and T500-Y41 have a very high occupancy in the BA.1.1 and BA.2 complexes but are weaker in BA.1 and BA.3. These interactions play a significant stabilizing role in both Wu-Hu-1 RBD and Omicron RBD complexes. At the same time, the important stable interactions Y449-D38 and Y489-Y83 and Y495-K353 that are very stable in the Wu-Hu-1 RBD complex are less stable in the Omicron RBD complexes (Table S2). The reduced occupancy of Y449-D38 interactions is partly due to the involvement of D38 in a salt bridge with R498 and hydrogen bonding with S496. Consistent with the previous studies [59], we found that minor differences in the RBD-ACE2 interfaces compositions (G446S and G496S in BA.1 and G446, G496 in BA.2) do not seem to significantly change the dynamics and stability of the intermolecular interactions in these positions (Figures 3,4). Structural studies proposed that the absence of the G496S mutation can be related to the enhanced affinity of BA.2 RBD [38] as S496 may affect the local interaction networks at the binding interface. According to the ensemble-based statistical analysis of the pairwise contacts in these systems, the interactions mediated by specific positions G496/S496 are rather similar and small binding energy differences between the Omicron subvariants may arise from cumulative contributions distributed over many interface residues.

### 2.3. Binding Free Energy Analysis and Electrostatic Interaction Potentials in the of the Omicron RBD-hACE2 Complexes

Using the conformational equilibrium ensembles obtained from 10 independent MD simulations, we computed the binding free energies for the Omicron RBD BA.1, BA.1.1, BA.2 and BA.3 complexes with hACE2 the MM-PBSA method [81] using the pipeline tool Calculation of Free Energy (CaFE) implemented as VMD plugin [82]. The binding free energy changes showed a fairly gradual but small improvement going from BA.1 to BA.1.1 and BA.2 RBD complexes (Table 3). At the same time, we found that the binding free energy changes for Omicron BA.3 complex are similar to the BA.1 variants.

**Table 3.** MM-PBSA binding free energies for the Omicron RBD-hACE2 complexes.*

| Systems | MM-PBSA Average Binding Free Energy (kcal/mol) | Standard Deviation (kcal/mol) | Experimental Binding Affinity |
| --- | --- | --- | --- |
| RBD Omicron BA.1-hACE2 | -48.785 | 6.345 | 19.5 nM |
| RBD Omicron BA.1.1-hACE2 | -50.565 | 7.235 | 5.9 nM |
| RBD Omicron BA.2-hACE2 | -53.125 | 6.454 | 10.0 nM |
| RBD Omicron BA.3-hACE2 | -49.778 | 6.976 | 22.1 nM |

\*The experimental binding affinities are from [38].

We also performed another binding free energy analysis using a contact-based predictor of binding affinity Prodigy [83], where computations of the binding affinity scores were averaged over 1,000 simulation samples from MD trajectories. (Table 4). In agreement with the MM-PBSA analysis, the results revealed a stronger binding affinity for the Omicron RBD BA.2 and BA.1.1 variants and somewhat weaker and similar binding free energies for the RBD BA.1 and BA.3 RBD complexes. The analysis of the interfacial inter-residue contacts revealed the larger number of stable interactions mediated by charged residues as well as the improved nonpolar interactions for the BA.1.1 and BA.2 complexes (Table 4). These observations indicated that the electrostatic interactions may be stronger in these complexes and together with the hydrophobic contacts drive the binding energetics of the Omicron RBD complexes. Overall, the predicted binding affinities reproduced the experimental data [38], but slightly favored BA.2 variant complex over BA.1.1 (Tables 3,4).

**Table 4.** The analysis of the interfacial residue-residue contacts and ensemble-averaged PRODIGY-based binding free energies for the Omicron RBD-hACE2 complexes.*

| Interactions | RBD Omicron BA.1-hACE2 | RBD Omicron BA.1.1-hACE2 | RBD Omicron BA.2-hACE2 | RBD Omicron BA.3-hACE2 |
| --- | --- | --- | --- | --- |
| Charged-charged | 7 | 8 | 9 | 6 |
| Charged-polar | 7 | 9 | 8 | 5 |
| Charged-apolar | 18 | 20 | 20 | 18 |
| Polar-polar | 5 | 5 | 6 | 6 |
| Polar-apolar | 17 | 16 | 20 | 18 |
| Apolar-apolar | 11 | 12 | 16 | 11 |
| $\Delta G$ computed(kcal/mol) | -11.2 | -11.8 | -12.3 | -11.2 |
| Kd (nM) experiment | 19.5 nM | 5.9 nM | 10.0 nM | 22.1 nM |

\*The experimental binding affinities are from [38].

The role of electrostatic interactions in stabilization of the Omicron RBD-ACE2 complexes has been extensively explored in previous computational studies [59-63]. To provide a detailed analysis of the electrostatic interactions between the Omicron RBD variants and hACE2, we employed the equilibrium ensembles obtained from MD simulations. The electrostatic interaction potentials are computed for the averaged RBD-hACE2 conformations for by solving Poisson Boltzmann equation using the APBS-PDB2PQR software [84,85] based on the Adaptive Poisson–Boltzmann Solver (APBS) [84] and visualized using the VMD visualization tool [86]. Using this approach, we first calculated the electrostatic potential on the surface of the cryo-EM SARS-CoV-2 S trimer structures for the Wu-Hu-1 closed and open states (Figure 5A,B), Omicron BA.1 variant (closed and open states) (Figure 5C,D), Omicron BA.2 (Figure 5E) and Omicron BA.3 variants (Figure 5F). Consistent with other studies [61], we found that the Wu-Hu-1 S trimers are characterized by the most negatively charged electrostatic surface in the S2 subdomain and S1/S2 regions (Figure 5A,B). where the electrostatic surface of the S1/S2 is mainly negatively charged and the S-NTD and S-RBD regions displayed a very moderately positively charged distribution.

**Figure 5.**
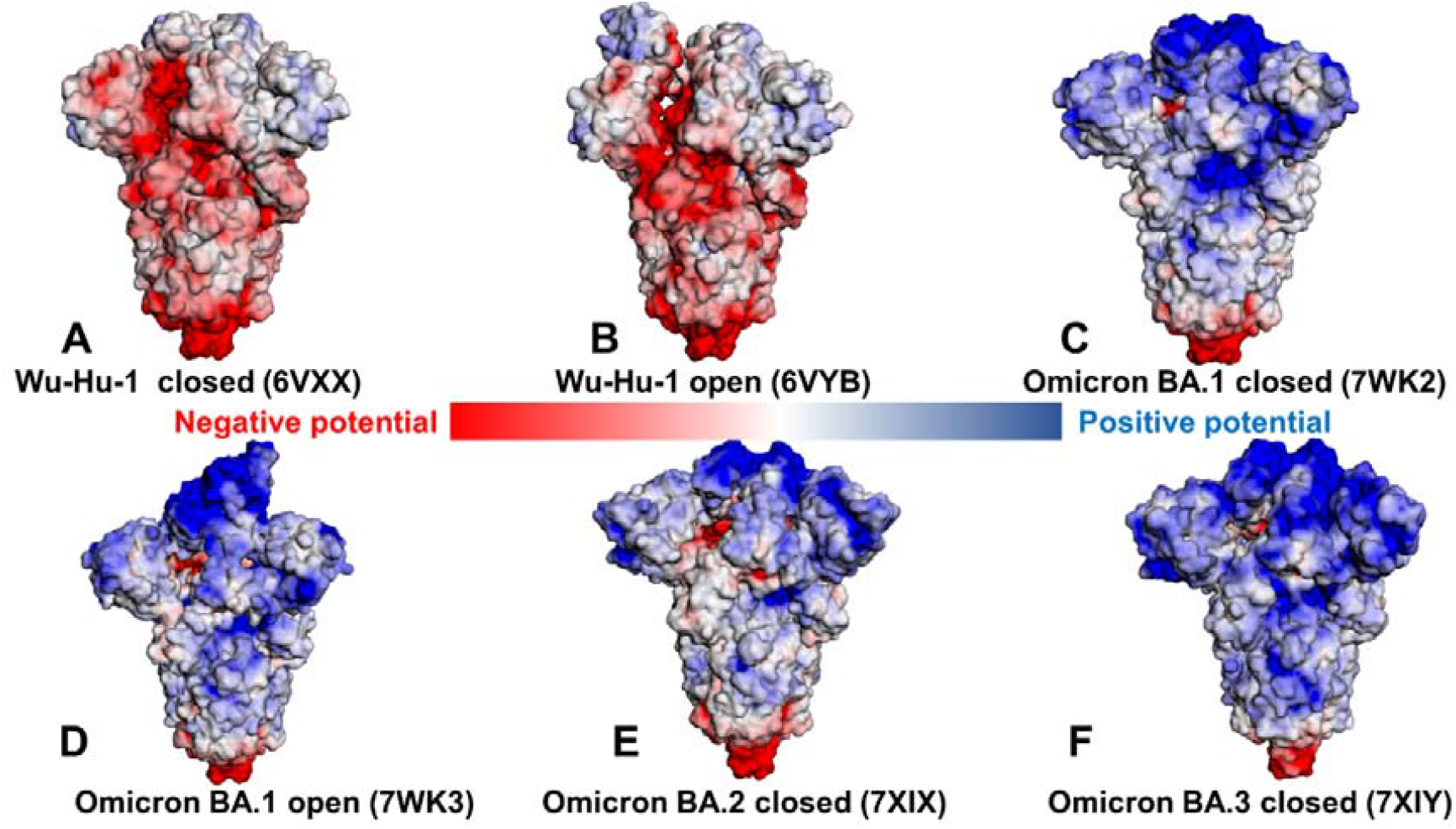
The distribution of the electrostatic potentials calculated with the APBS method [84,85] on the molecular surface of trimeric S proteins of Wuhan strain (Wu-Hu-1) closed state, pdb id 6VXX (A), S-Wu-Hu-1 trimer in the open state, pdb id 6VYB (B), S-Omicron BA.1 closed trimer, pdb id 7WK2 (C), S-Omicron BA.1 open 1-up trimer, pdb id 7WK3 (D), S-Omicron BA.2 closed trimer, pdb id 7XIX (E), and S-Omicron BA.3 closed trimer, pdb id 7XIY (F). The color scale of the electrostatic potential surface is in units of kT/e at T = 37°C. Electro-positively and electronegatively charged areas are colored in blue and red, respectively. Neutral residues are in white.

The analysis of the closed and open states for the Wu-Hu-1 S trimers particularly highlighted variations in the electrostatic distribution of the RBD regions, pointing to the absence of strong positively charged regions (Figure 5A,B). In a marked contrast, the Omicron S trimer has a strongly positive S1 subdomain, clearly displaying a strong positively charged surface in the S-NTD and S-RBD regions (Figure 5C,D). Indeed, a number of positively charged sites in the Omicron RBD regions emerged through mutational changes N440K, T478K, Q493R, Q498R and Y505H. A comparison of the electrostatic potential surfaces for the Omicron variants (Figure 5C-F) illustrated that rapid change in the distribution between Wu-Hu-1 and Omicron BA.1 may be contrasted with a more gradual incremental evolution among BA.1, BA.1.1, BA.2 and BA.3. The overall differences in the electrostatic potential surfaces for the Omicron subvariants are relatively moderate, but we observed a greater accumulation of the positively charged regions in the BA.2 and BA.3 subvariants (Figure 5E,F). These findings are consistent with related investigations [59-61] that emphasized a decisive role of the electrostatic interactions in driving binding signatures of the S protein, particularly suggesting that ACE2 binding can be modulated via electrostatic contributions and antibodies that are escaped by Omicron variants may have positive electrostatic interacting surfaces [61]. In general, despite mutational differences between BA.1, BA.1.1, BA.2 and BA.3 lineages, the electrostatic potential surfaces for the respective S trimer structures are similar, showing a strong positive electrostatic surface in the S-RBD regions which interface with the negatively charged ACE2 receptor.

We then proceeded to compute the electrostatic potential surfaces of the RBD-hACE2 complexes for the ensemble-averaged structures across all Omicron subvariants (Figure 6). Similarly, we found a stronger positively charged electrostatic distribution on the Omicron RBDs (Figure 6B-E) as compared to the original strain (Figure 6A). The surface of hACE2 is enriched by the negatively charged residues, which is particularly apparent in the H1 helix of hACE2 involved in the key interactions with the RBD regions. The electrostatic potentials on the RBD surfaces for the Omicron RBD-hACE2 complexes are mainly positive with variable charge distributions, showing relatively moderate changes in the overall surface distribution between subvariants. Nonetheless, we noticed a stronger positively charged potential in the Omicron RBD BA.1.1 (Figure 6C) showing positive densities in the RBM-interacting regions and also in distal RBD regions not directly involved in the contacts with the hACE2. According to structural studies [38], R346K in BA.1.1 RBD enhances the interaction with hACE2 through long-range alterations by allosterically mediating stronger electrostatic interactions of R493 with E35 and D38 sites. These subtle changes in the RBD interaction networks can be reflected in the increased electrostatic potential of the RBD BA.1.1 subvariant bound to ACE2 (Figure 6C). However, it is worth pointing out that the corresponding changes in the electrostatic potentials between studied Omicron RBD variants are relatively minor. Using DelPhiPKa protonation and pKa calculations [88,89], we also computed the electrostatic polar energies and desolvation energies for the RBD residues based on the equilibrium structures of the Omicron RBD-hACE2 complexes (Tables S3-S6). We particularly examined the electrostatic polar energies for individual residues in their protonated and neutral states, focusing specifically on key RBD binding residues R493, R498 and Y501 involved in the network of favorable interactions across all Omicron RBD-hACE2 complexes. The results showed relatively similar and favorable electrostatic polar energies for these residues in the protonated sate in the BA.1 and BA.1.1 variants (Tables S3,S4) but also revealed the stronger electrostatic interactions in these positions for the Omicron BA.2 complex (Table S5). Hence, computations of the electrostatic potential surfaces confirmed that the Omicron RBD variants have a stronger positive electrostatic potential due to the substitutions to basic residues including N440K, T478K, Q493R and Q498R, whereas E484A can reduce a negative charge on the RBD. The binding interface of ACE2 in the S-RBD Omicron complexes showed a significant negative electrostatic potential, providing the improved complementarity of the intermolecular surfaces as compared to the original Wu-Hu-1 strain.

**Figure 6.**
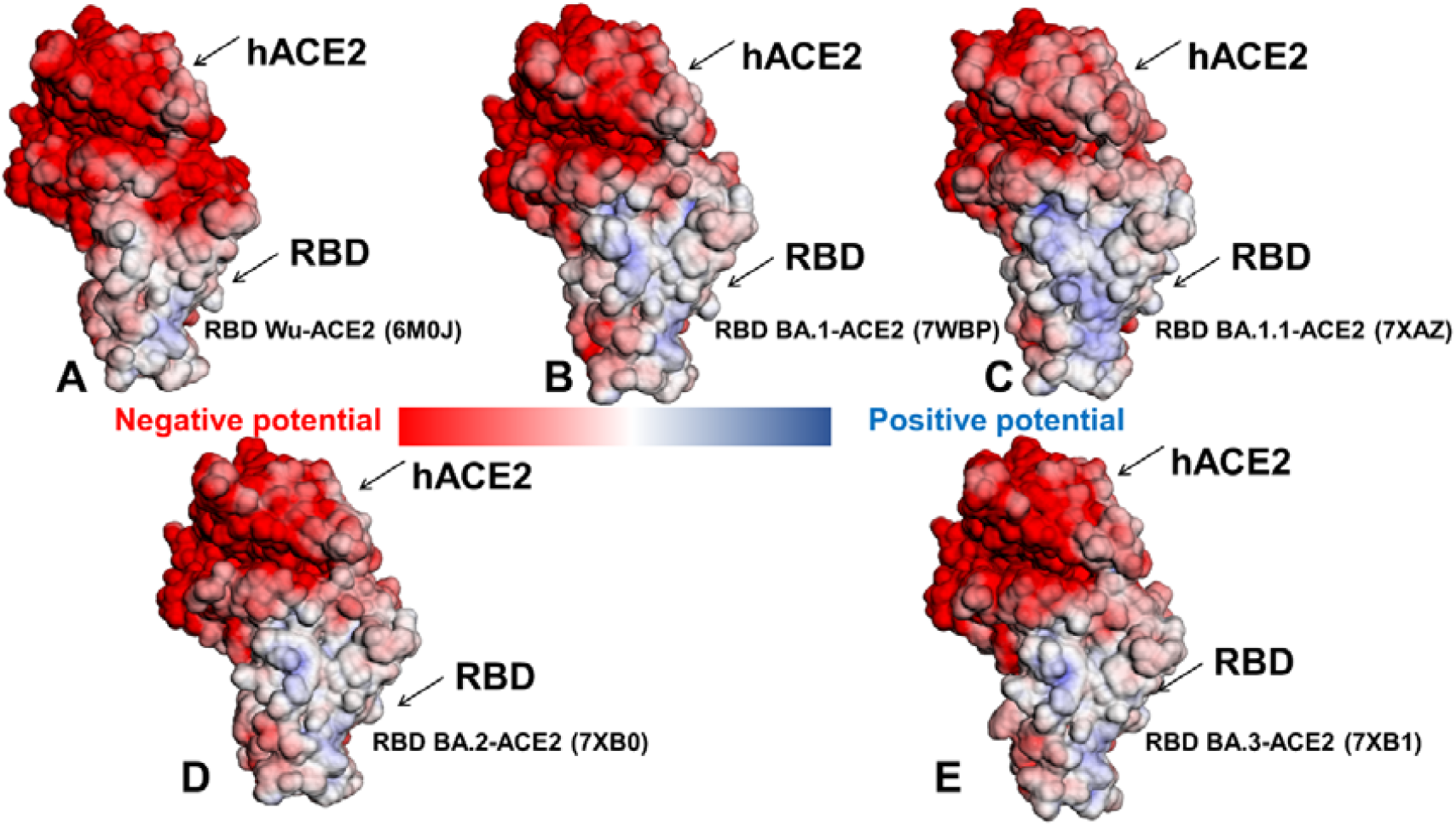
The distribution of the electrostatic potentials calculated with the APBS method [84,85] on the molecular surface of the S-RBD Wu-Hu-1 complex with hACE2, pdb id 6M0J (A), S-RBD Omicron BA.1 complex with hACE2, pdb id 7WBP (B), S-RBD Omicron BA.1.1 complex with hACE2, pdb id 7XAZ (C), S-RBD Omicron BA.2 complex with hACE2, pdb id 7XB0 (D), and S-RBD Omicron BA.3 complex with hACE2, pdb id 7XB1 (E). The color scale of the electrostatic potential surface is in units of kT/e at T = 37°C. Electro-positively and electronegatively charged areas are colored in blue and red, respectively. Neutral residues are in white.

The analysis of the stable salt bridges and hydrogen bonds based on simulations also demonstrated a considerable increase in stabilization of key interactions mediated by R493 and R498 (R493-E35, R493-D38, R498-D38) in the BA.1.1 and BA.2 complexes (R493-E35, R493-D38) (Table S2). We found that Q493R and Q498R sites form new and extremely stable salt bridges with the electronegative E35 and D38 residues of hACE2, which is likely one of the main contributors of the enhanced binding affinity of the Omicron RBD-hACE2 complexes. The stable R493-E35 and R498-D38 interactions are shared among all Omicron subvariants (Table S1). In addition, stable R493-D38 interactions are characteristic of the Omicron RBD BA.1.1 complex, but these additional contacts are absent in other complexes. A significant conservation of stable R498-Y41, T500-Y41, and Y501-Y41 interactions in MD simulations is also observed across all Omicron RBD complexes (Table S1). Together, the results of MD simulations, ensemble-based contact analysis and mapping of the electrostatic potentials consistently pointed to the critical roles of R493, R498 and particularly Y501 in mediating strong binding interactions for the Omicron RBD complexes. Our results support the previous studies [90-92] in asserting functional importance of these interaction hotspots in the ACE2 binding. To summarize, the presented integrated analysis reinforced the notion that the enhanced positively charged potential of the RBD regions in the Omicron RBD variants could contribute to stronger binding with the negatively charged host receptor ACE2. At the same time, the revealed minor differences in the electrostatic potentials between Omicron RBD subvariants suggested a more subtle mechanism for the enhanced binding of the BA.1.1 and BA.2 which is likely associated with small cumulative changes distributed over many RBD residues.

### 2.4. Ensemble-Based Mutational Sensitivity Analysis Identifies Key Structural Stability and Binding Affinity Hotspots in the SARS-CoV-2 RBD Complexes with ACE2

Using the conformational ensembles of the Omicron RBD variant complexes we performed a systematic mutational scanning of the interfacial RBD residues. In silico mutational scanning was done using BeAtMuSiC approach [93-95]. This approach utilizes statistical potentials and allows for robust predictions of the mutation-induced changes of the binding interactions and the stability of the complex. We enhanced BeAtMuSiC approach by invoking ensemble-based computations of the binding free energy changes caused by mutations. In our adaptation of this method, the binding free energy ΔΔG changes were evaluated by averaging the results of computations over 1,000 samples from MD simulation trajectories. To provide a systematic comparison, we constructed mutational heatmaps for the RBD interface residues in each of the studied Omicron RBD-hACE2 complexes (Figure 7). Consistent with deep mutagenesis experiments [68,96-99], the strongest stability and binding energy hotspots correspond to hydrophobic residues F456, Y489 and Y501 that play a decisive role in binding for all Omicron complexes (Figure 7). Mutational heatmaps clearly showed that all substitutions in these key interfacial positions can incur a consistent and considerable loss in the stability and binding affinity with ACE2. Interestingly, these positions are distributed across both major patches of the binding interface. These observations are also consistent with the importance of ACE2 hotspots K31 and H34 in the middle of the interface forming an extensive interaction network with Y453, L455, F456, and Q493 residues. The common energetic hotspots Y453, F456, Y489 and Y501 found in the mutational scanning also emerged as critical stability and binding hotspots in deep mutagenesis studies [68,96-99]. It is worth stressing that these universal hotspots include conserved residue Y489 and homologues position F456 along with the variable hotspot N501Y. Our analysis confirmed that theses RBD residues are universally important for binding in all Omicron variants and act as molecular anchors of the RBD stability and binding affinity (Figure 7). The results showed that the Omicron RBD preserves key stability centers Y453, F456 and Y489 while acquiring the additional binding affinity and sensitivity primarily through the N501Y mutated site. This is consistent with the experimental data showing that N501Y mutation alone can improve the binding affinity by 6-fold [100]. Although structural and dynamic similarities between Omicron RBD complexes yielded fairly similar energetic heatmaps, there are a number of notable differences that provide an insight into binding mechanisms and reveal functional roles of RBD residues in binding and stability. The mutational heatmap for the Omicron RBD BA.1 revealed an important role of Y453, F456, Y489, Q493R, T500, N501Y and Y505H (Figure 7A). A non-canonical π–π stacking interaction between Y501 and ACE2 residue Y41 and additional interactions with ACE2 binding hotspot K353 provides a considerable stabilization. As a result, all modifications in the Y501 position of the Omicron RBD BA.1 complex produced significant destabilization changes (Figure 7A). A comparison of mutational heatmaps for RBD BA.1 (Figure 7A) and RBD BA.1.1 complexes (Figure 7B) are generally similar, highlighting the presence of several binding hotspot clusters. One group of mutation-sensitive residues includes Y453, L455 and F456 while other hotspots corresponded to Y489 and group of residues centered around Y501 (T500, G502, and H505). Another relatively sensitive position is Q493R. In the Omicron RBD BA.1 complex, R493 forms new salt bridges with E35, while in the Omicron RBD BA.1.1 the interfacial contacts of R493 with K31, H34, E35 and D38 in result in a denser electrostatic intermolecular network Mutational scanning in R493 showed larger destabilization changes in the BA.1.1 complex as compared to the RBD BA.1 complex. For instance, the binding free energy changes for R493A (ΔΔG = 1.58 kcal/mol), R493D (ΔΔG = 1.58 kcal/mol), R493K (ΔΔG = 0.65 kcal/mol) in BA.1 complex can be contrasted with the corresponding changes of ΔΔG = 1.85 kcal/mol, ΔΔG = 1.74 kcal/mol and ΔΔG = 1.19 kcal/mol respectively in the BA.1.1 complex (Figure 7A,B). In both BA.1 and BA.1.1 complexes Q498R makes new contacts with D38 and Q42, G496S and Y505H form new hydrogen bond with K353. As a result, a consistent destabilization pattern of the binding free energy changes was seen for R493, S496, and R498 positions of the Omicron RBD BA.1 and BA.1.1 complexes (Figure 7A,B).

**Figure 7.**
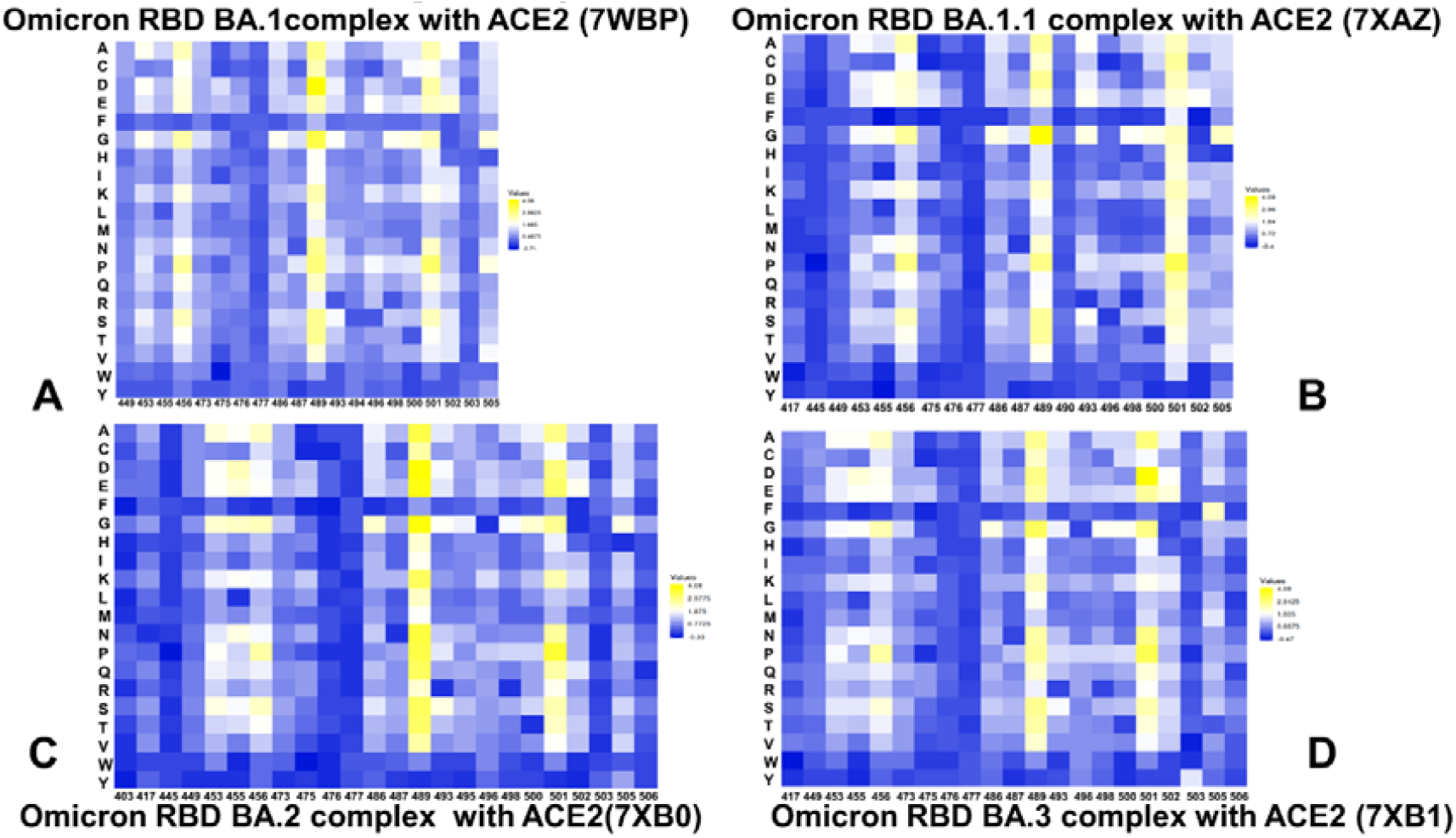
Ensemble-based dynamic mutational profiling of the RBD intermolecular interfaces in the Omicron RBD-hACE2 complexes. The mutational scanning heatmaps are shown for the interfacial RBD residues in the Omicron RBD BA.1-hACE2 (A), Omicron RBD BA.1.1-hACE2 (B), Omicron RBD BA.2-hACE2 (C), and Omicron RBD BA.3-hACE2 complexes (D). The heatmaps show the computed binding free energy changes for 20 single mutations of the interfacial positions. The cells in the heatmap are colored using a 3-colored scale – from blue to white and to yellow, with yellow indicating the largest unfavorable effect on binding. The standard errors of the mean for binding free energy changes were based on MD trajectories and selected samples (a total of 1,000 samples) are within ∼ 0.08-0.2 kcal/mol using averages over 10 different MD trajectories.

Mutational scanning of the RBD residues in the Omicron RBD BA.2 complex highlighted a larger binding interface, in which the mutation-induced destabilization is distributed over more residues (Figure 7C). The large destabilization changes become more pronounced for Y453, L455 and F456 while also revealing highly sensitive positions for F486, N487, Y489, R493, T500, Y501, and H505 (Figure 7C). The difference in the RBD-ACE2 interface for the BA.1 and BA.2 variants are the two residues located at residue positions 446 and 496, which are S446 and S496 for BA.1 and G446 and G496 for BA.2. While S496 makes interaction contacts with D38 and K353, G496 makes contacts with D38 in the BA.2 complex. Interestingly, mutational scanning in S496 position in the RBD BA.1 and BA.1.1 complexes revealed moderate destabilization changes in the range ΔΔG = 0.3 −1.5 kcal/mol, where S496G mutation is destabilizing with ΔΔG =0.63 kcal/mol for the BA.1 complex and ΔΔG =0.48 kcal/mol for the BA.1.1 complex (Figure 7C). At the same time, mutations of G496 in the BA.2 complex showed larger destabilization changes than corresponding modifications of S496 in the BA.1.1 complex. In particular, the reversed G496S mutation is more destabilizing in the BA.2 complex with ΔΔG =0.88 kcal/mol than the respective S496G mutations in the BA.1 and BA.1.1 variants. Based on structural observations, it was suggested that the S496G site mutation could contribute to ∼ 3.6-fold affinity increase, whereas the G496S mutation in BA.2 RBD may induce ∼ 1.5-fold decrease in hACE2 binding affinity [38]. Our analysis showed that both S496 and G496 sites make favorable interactions in their respective RBD Omicron complexes, but the contribution of G496 in the BA.2 complex may be moderately more significant than the binding effect of S496 in the BA.1. Hence, our results are consistent with the experimental data [38] and provide a plausible rationale for subtle differential effects induced by S496/G496 in the Omicron complexes.

The destabilization changes induced by mutations in Y453, L455 and F456 positions in the BA.2 and BA.3 complexes are greater than in the other Omicron RBD complexes. Another patch of binding energy hotspots corresponds to F486, N487 and Y489 residues (Figure 7C,D). In this case, modifications of F486 lead to larger free energy changes in the BA.2 and BA.3 complexes. The extended patch of binding interface residues in the RBD BA.2 and BA.3 complexes (R493, Y495, G496, R498, T500, Y501, G502 and H505) contributes significantly to binding. Most notably mutational scanning of R493, R498 and Y501 positions in the RBD BA.2 and BA.3 complexes resulted in considerable destabilization free energies that are larger than in the BA.1 and BA.1.1 complexes(Figure 7). These findings suggested that the stronger binding affinity of the BA.2 complex may be associated with the larger binding interface that allows for the greater cumulative contribution of the RBD binding residues. At the same time, we observed that the binding energy hotspots (Y453, L455, F456, Y489, R493, R498, Y501) are shared between BA.2 and BA.3 complexes, yielding the bulk of the binding energy (Figure 7C,D). Our results indicated that Y501 position is the most critical binding affinity hotspot in the Omicron RBD complexes with ACE2. Moreover, a tight cluster of binding affinity hotspots in this region formed by Y501, R498, S496/G496 and H505 residues make the dominant contribution to the binding affinity (Figure 4E). These binding energy hotspots are packed together and collectively form a dense network of the hydrophobic and polar interactions with ACE2. By combining insights obtained from binding affinity analysis and mutational scanning, we argue that the enhanced binding affinity of the BA.2 complex may be determined by the strengthened inter-molecular electrostatic interactions mediated by R493, R498 and Y501 as well as cumulative contributions distributed over larger binding interface with the hACE2 receptor.

To compare the mutation-induced changes in the protein stability and binding interactions of the Omicron RBD-hACE2 complexes, we also performed a systematic mutational scanning of all RBD residues and evaluated the resulting folding free energy changes and binding free energy changes using enhanced BeAtMuSiC approach (Figure 8A-D). The density distributions of mutation-induced folding free energy changes in the Omicron RBD complexes are generally similar and are characterized by single peak around ΔΔG = 0.5 kcal/mol accompanied by a fairly long tail extending to ΔΔG = 2.5 kcal/mol (Figure 8A-D). The overall distributions of the binding free energies displayed a distinct shape featuring multiple peaks over broader range of ΔΔG values, where one of the peaks corresponds to ΔΔG = 0.5 kcal/mol but shallow peaks observed at ΔΔG −1.5 −2 kcal/mol. Notably, the distribution of folding free energy changes is entirely characterized by the destabilization stability changes (ΔΔG > 0.0) but the average free energy values of ΔΔG ∼ 0.5 kcal/mol indicated relatively minor destabilization effects (Figure 8). The binding free energy changes for the RBD residues showed a very minor density for the negative ΔΔG values of −0.2 to - 0.5 kcal/mol associated with the improved binding, while the density profile indicated that mutations on average have a destabilization effect on binding (Figure 8A-D). The scatter plots of the free energy changes for all RBD residues in the Omicron RBD complexes and the original Wu-Hu-1 RBD complex showed a strong correlation (Figure 8E-H), reflecting significant structural, dynamic and energetic similarities between these systems. Nonetheless, these graphs indicated some minor dispersion where some mutations with relatively small ΔΔG =0.5-1.0 kcal/mol in the Wu-Hu-1 RBD complex produce considerably larger ΔΔG = 2.0-3.0 kcal/mol in the Omicron RBD complexes (Figure 8E-H).

**Figure 8.**
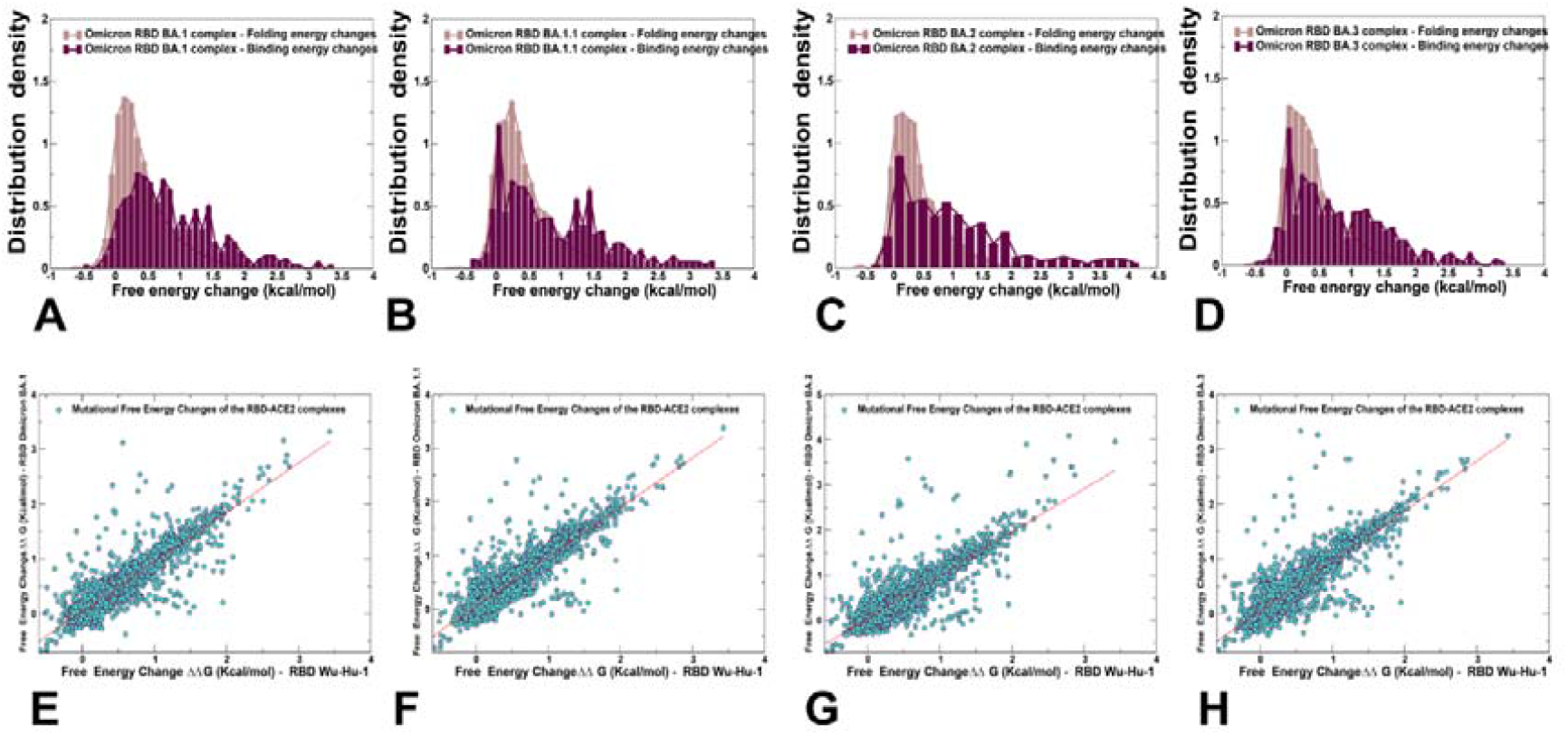
Ensemble-based distributions of the RBD folding free energy changes in the Omicron RBD-complexes based on mutational scanning of all RBD residues (in light brown bars) and biding energy changes based on mutational scanning of the intermolecular residues (in maroon bars). The density distributions are shown for the Omicron RBD BA.1-hACE2 (A), Omicron RBD BA.1.1-hACE2 (B), Omicron RBD BA.2-hACE2 (C), and Omicron RBD BA.3-hACE2 complexes (D). The 2D scatter plots of the free energy changes for the RBD residues in the Omicron RBD-hACE2 complexes. The scatter plot of the free energy changes from mutational scanning of the RBD residues in the reference RBD Wu-Hu-1 complex and the Omicron RBD BA.1 complex (E), RBD BA.1.1 complex (F), RBD BA.2 complex (G), and RBD BA.3 complex (H). The data points are shown in light-brown colored circles.

In general, this analysis suggests that the evolutionary paths for significant improvements in the binding affinity of the Omicron RBD variants with hACE2 are relatively narrow and require balancing between various fitness tradeoffs of preserving RBD stability, maintaining binding to ACE2, and allowing for immune evasion [101]. These factors may severely limit the “ evolutionary opportunities” for the virus to adapt new mutations that markedly improve ACE2 binding affinity without compromising immune evasion and stability. These arguments are consistent with the recent realization that evolutionary pressure invokes a complex interplay of thermodynamic factors between mutations that retain or marginally improve affinity for the ACE2 with other RBD modifications facilitating immune escape. Moreover, the emerging consensus proposed that immune evasion may be a primary driver of Omicron evolution that sacrifices some ACE2 affinity enhancement substitutions to optimize immune-escaping mutations [29-33].

### 2.5. Dynamic Modeling of the Residue Interaction Networks and Community Analysis of the Omicron RBD Complexes Detail Role of Binding Hotspots in Mediating Epistatic Interaction Effects

The evolutionary and functional studies [67-71] suggested that evolutionary windows for the Omicron variants could be enhanced through epistatic interactions between variant mutations and the broader mutational repertoire. It was suggested that weak epistasis for the Wu-Hu-1 original strain may become more pronounced for the Omicron variants as a potential virus mechanism to counter limited potential for continued evolution [67]. In this section, we employed the ensemble-based modeling of the residue interaction networks utilizing dynamics and coevolution [102-105] together with community decomposition analysis to propose a topology-based model to characterize non-additive effects between mutational sites in the Omicron RBD complexes. To characterize potential non-additive effects of Omicron mutations, we investigated the amino-acid network structures for each mutant and determined the topological quantities of these networks. In this approach, the dynamic fluctuations are mapped onto a graph with nodes representing residues and edges representing weights of the measured dynamic properties [102] and coevolutionary couplings between residues [103]. Our previous network modeling studies of SARS-CoV-2 S proteins showed that this approach can adequately capture the nature of allosteric interactions and communications in the S proteins [72-78].

Based on the dynamic inter-residue interaction networks, we performed community decomposition and characterized stable local interaction modules in which residues are densely interconnected through coupled interactions and dynamic correlations (Figure 9). A community-based model of allosteric interactions employed in this study is based on the notion that groups of residues that form local interacting communities are correlated and switch their conformational states cooperatively. In this model allosteric communications are transmitted through a cascade of stable local modules connected via inter-community bridges. Within the network analysis, we also characterized the distribution of clique communities. In network terms, a k-clique is defined a complete sub-graph of k nodes in which each pair of nodes is connected by an edge, reflecting strong interactions and dynamic coupling between every node in the clique with all other nodes that belong to the same clique. A collection of all interconnected k-cliques in a given network defines a k-clique community. We assumed that if the mutational sites of the Omicron RBD complexes fall into to a 3-clique structure we can associate this topological organization as a signature of potential non-additivity effects. In other words, mutational positions that provide an indispensable stabilizing anchor to multiple intermolecular RBD-ACE2 cliques are assumed to have a stronger non-additive effect on binding. Using topological network analysis and characterizing the distribution of stable 3-clique communities, we evaluated the role of RBD binding residues in mediating potential epistatic effects in the Omicron RBD complexes.

**Figure 9.**
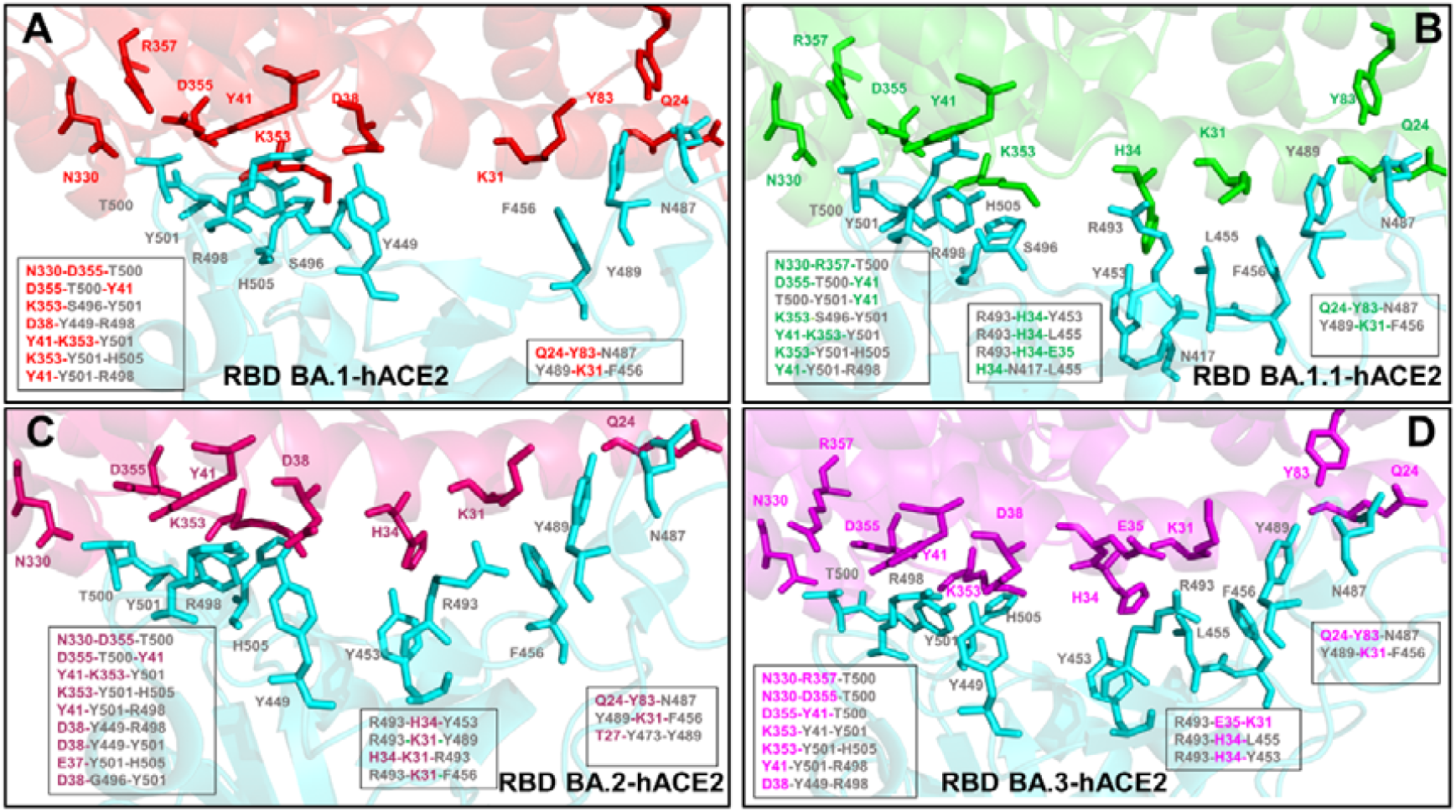
The network-based community analysis to infer potential epistatic relationships between S-RBD residues. The distributions of stable 3-clique communities formed at the binding interface of the Omicron RBD complexes with hACE2. Structural mapping and full annotation of the intermolecular cliques for the RBD BA.1-hACE2 complex. The RBD binding residues are shown in cyan sticks and the hACE2 binding residues are in red sticks (A). Structural mapping and annotation of the intermolecular cliques for the RBD BA.1.1-hACE2 complex. The RBD binding residues are shown in cyan sticks and the hACE2 binding residues are in green sticks (B). Structural mapping and full annotation of the intermolecular cliques for the RBD BA.2-hACE2 complex. The RBD binding residues are shown in cyan sticks and the hACE2 binding residues are in pink sticks (C). Structural mapping and full annotation of the intermolecular cliques for the RBD BA.3-hACE2 complex. The RBD binding residues are shown in cyan sticks and the hACE2 binding residues are in purple sticks (D).

The analysis of stable 3-clique communities formed at the binding interface of the Omicron RBD BA.1 complex showed a dense and lopsided asymmetric distribution with seven distinct cliques formed in the vicinity of Y501 site (Figure 9A). These RBD-ACE2 interfacial cliques include N330-D355-T500, D355-T500-Y41, K353-S496-Y501, D38-Y449-R498, Y41-K353-Y501, K353-Y501-H505, and Y41-Y501-R498 clusters (Figure 9A). Strikingly, the majority of these stable clusters are anchored by Y501 position. Moreover, the clique distribution showed a strong interaction coupling between Y501 and R498 in binding Y41 on ACE2, and R498 coupled with Y449 in stabilizing binding contacts with another ACE2 hotspot D38. Additionally, Y501 appeared to be strongly linked with S496, Y449, and T500 in forming multi-residues interfacial cliques in the Omicron RBD BA.1 complex (Figure 9A). These observations are in excellent agreement with illuminating functional studies that provided compelling evidence of epistatic shifts in the RBD driven by the N501Y mutation [68]. In these experiments, using deep mutational scanning it was discovered that the RBD sites exhibit notable epistatic shifts due to coupling to N501Y and the largest epistatic shift in mutational effects is associated with non-additive contribution of Q498R and N501Y, followed by less pronounced epistatic shifts at sites 491 to 496 at the ACE2 contact surface [68]. Consistent with these experiments, our network model showed that not only structurally proximal R498 and Y501 are strongly coupled and could therefore exhibit epistatic couplings, but also several other RBD residues S496, T500 and H505 form a group of sites that are highly correlated with Y501 and ACE2 binding residues. Hence, the clique-based network model can identify highly correlated and potentially non-additive mutational sites in the Omicron RBD complexes and distinguish them from other mutational sites that are less likely to experience epistatic shifts.

While the mutational scanning maps of the Omicron RBD interfacial residues are quite similar, the differences in the distribution of the dynamic network cliques between variants are quite apparent and often more instructive. For the Omicron RBD BA.1.1 complex, the clique distribution is considerably broader and covered distinct patches of the binding interface with ACE2 (Figure 9B). In this complex, the dominant contribution of stable cliques near Y501 is preserved, and the clusters N330-D355-T500, D355-T500-Y41, T500-Y501-Y41, K353-S496-Y501, Y41-K353-Y501, K353-Y501-H505, and Y41-Y501-R498 emerged as conserved signatures of the binding interfaces (Figure 9B). Strikingly, for the BA.1.1 complex, we observed several stable dynamic cliques in the middle segment of the binding interface, namely R493-H34-Y453, R493-H34-L455, R493-H3-E35, and H34-N417-L455. A group of stable cliques was also found in the adjacent patch and included Q24-Y83-N487 and Y489-K31-F456 (Figure 9B). In the Omicron RBD BA.2 complex, we similarly found a dense and patch-connected distribution of the interfacial cliques (Figure 9C). The large proportion of the interfacial 3-cliques is consolidated in the patch anchored by the Y501 site. A total of nine different cliques in this region was detected (N330-D355-T500, D355-T500-Y41, Y41-K353-Y501, K353-Y501-H505, Y41-Y501-R498, D38-Y449-R498, D38-Y449-Y501, E37-Y501-H505, and D38-G496-Y501) (Figure 9C). Notably, the reversal of G446S and G496S in BA.1/BA.1.1 to G446 and G496 in BA.2 resulted in the loss of the K353-S496-Y501 interfacial clique but was compensated by the emergence of several additional cliques E37-Y501-H505, and D38-G496-Y501 that are not present in BA.1.1. Overall, we found the largest number of three-residues cliques in the Omicron RBD BA.2 complex which reflects the stability and strength of the intermolecular interactions for this subvariant. These observations also suggested that the extent of non-additive epistatic contributions in the Omicron RBD BA.2 complex can be stronger than in the other subvariants (Figure 9C). It is worth emphasizing that most of these cliques are anchored by the Y501 site that emerged as a key epistatic hub that can mediate strong couplings with other RBD sits and ACE2 binding residues. The RBD residues Y449, G496, R498, and T505 are dynamically coupled with Y501 in the various binding interfacial cliques, revealing a dominant role of Y501 as a dynamic network mediator of non-additive contributions to the binding affinity of the BA.2 variant. These observations are in excellent agreement with the deep mutational scanning which showed that the major sites exhibiting epistatic shifts in the presence of Y501 include R498, G446, G496, Y499 residues [68]. In addition, for the RBD BA.2 complex, we detected a number of 3-cliques in the middle patch of the interface (R493-H34-Y453, R493-K31-Y489, H34-K31-R493, and R493-K31-F456) and only a small number of cliques on the other side of the interface (Q24-Y83-N487, Y489-K31-F456, and T27-Y473-Y489) (Figure 9C). These results highlighted the role of R493 position in anchoring multiple interaction clusters with the ACE2 residues, also indicating some level of dynamic coupling with Y453 and Y489 residues. The conserved Y453 and Y489 positions are involved in favorable hydrophobic interactions and dynamic coupling with the interactions mediated by R493 suggest some level of cooperativity and synchronicity between these contributions. At the same time, these findings indicated that R493 is not strongly dynamically coupled with other Omicron RBD mutational sites and consequently may be less important in mediating broad epistatic shifts which is consistent with the experimental data [68].

The distribution of stable cliques in the Omicron RBD BA.3 complex is generally similar to that of RBD BA.2 complex, featuring an extensive network of dynamic cliques in the patch anchored by Y501 (N330-D355-T500, N330-R357-T500, D355-T500-Y41, Y41-K353-Y501, K353-Y501-H505, Y41-Y501-R498, D38-Y449-R498) (Figure 9D). In comparison with the BA.2 complex, we did not detect stable cliques that involve G496 position on RBD, suggesting that the interactions mediated by G496 with D38 and Y501 are very dynamic and coupling between Y501 and G496 could be transient and fairly weak to warrant appreciable epistatic shifts. The spectrum of 3-cliques in the middle patch is determined by R493 and include R493-E35-K31, R493-H34-L455 and R493-H34-Y453 clusters (Figure). Notably, for both BA.2 and BA.3 complexes, we found a strong dynamic coupling between R493, Y453 and L455 forming a dense and interconnected network with K31, H34, and E35 binding hotspot on ACE2 (Figure 9D). This is consistent and is partly reflected in the mutational scanning maps for these Omicron complexes that identified these RBD positions as important binding hotspots (Figure 7). The destabilization effect of mutations in these sites is more significant than in the BA.1 and BA.1.1 complexes, indicating a broadening of the binding hotspot residues in the BA.2 and BA.3 subvariants.

To summarize, the network-based community analysis provided an additional insight to the mutational scanning data showing that mutational changes in these three positions may be coupled and lead to non-additive negative epistatic effects. However, importantly, these effects are secondary as the major non-additive contributions are likely to arise from presence of large number of stable cliques mediated by Y501 position. Our findings suggested that the extent of non-additive contributions to the binding affinity may be greater for the Omicron BA.1.1 and particularly BA.2 complex that displayed the strongest binding affinity among the examined Omicron subvariants.

### 2.6. Reversed Allosteric Communication Analysis of the Omicron RBD Complexes Identifies Conserved Allosteric Binding Site and Suggests that Allosteric Hotspots Anchor Allosteric Pockets in the RBD

Here, we employed and adapted the reversed allosteric communication approach based on the network modeling of the Omicron RBD-hACE2 complexes to compute global topological metrics that characterize the network modularity and inter-community connectivity. We propose that by modeling allosteric communication networks and identifying allosteric mediating hotspots using network parameters, we can characterize the distribution and topology of allosteric pockets on the RBD. We will show that via this reversed allosteric communication approach, we can map the experimentally known allosteric binding site in the RBD core which is distal from the RBD-hACE2 binding interface. The reversed allosteric communication approach is based on the premise that allosteric signaling in proteins is bidirectional and can propagate from an allosteric to orthosteric site and vice versa [106-109]. Some of the reversed allosteric communication approaches proposed by our laboratory and others are rooted in the dynamic network-based models of inter-residue interaction [110-113]. A more integrated computational and experimental strategy exploited the reversed allosteric communication concepts to combine MD simulations and Markov state models (MSM) for characterization of binding shifts in the protein ensembles and identification of previously unknown cryptic allosteric sites [108,114]. In the current study, we constructed dynamic inter-residue interaction networks and computed the short path residue centrality (SPC) to analyze modularity and allosteric communications. The SPC distributions reflect the extent of residue connectivity in the interaction networks and allow for characterization of the mediating clusters in the complexes. By identifying local residue clusters that correspond to the major peaks of the SPC distributions, we infer positions of regions harboring potential allosteric hotspot points that mediate long-range interactions and communications in the Omicron RBD-hACE2 complexes (Figures 10,11). Using this network-based reconstruction of allosteric communications in the RBD complexes, we then dynamically map clusters of residues involved in mediating allosteric communications onto average conformational states of the Omicron RBD complexes (Figures 10,11). In our model, we hypothesized that these allosteric clusters would overlap with the allosteric binding sites and provide a robust reversed allosteric communication approach for characterization of allosteric pockets.

**Figure 10.**
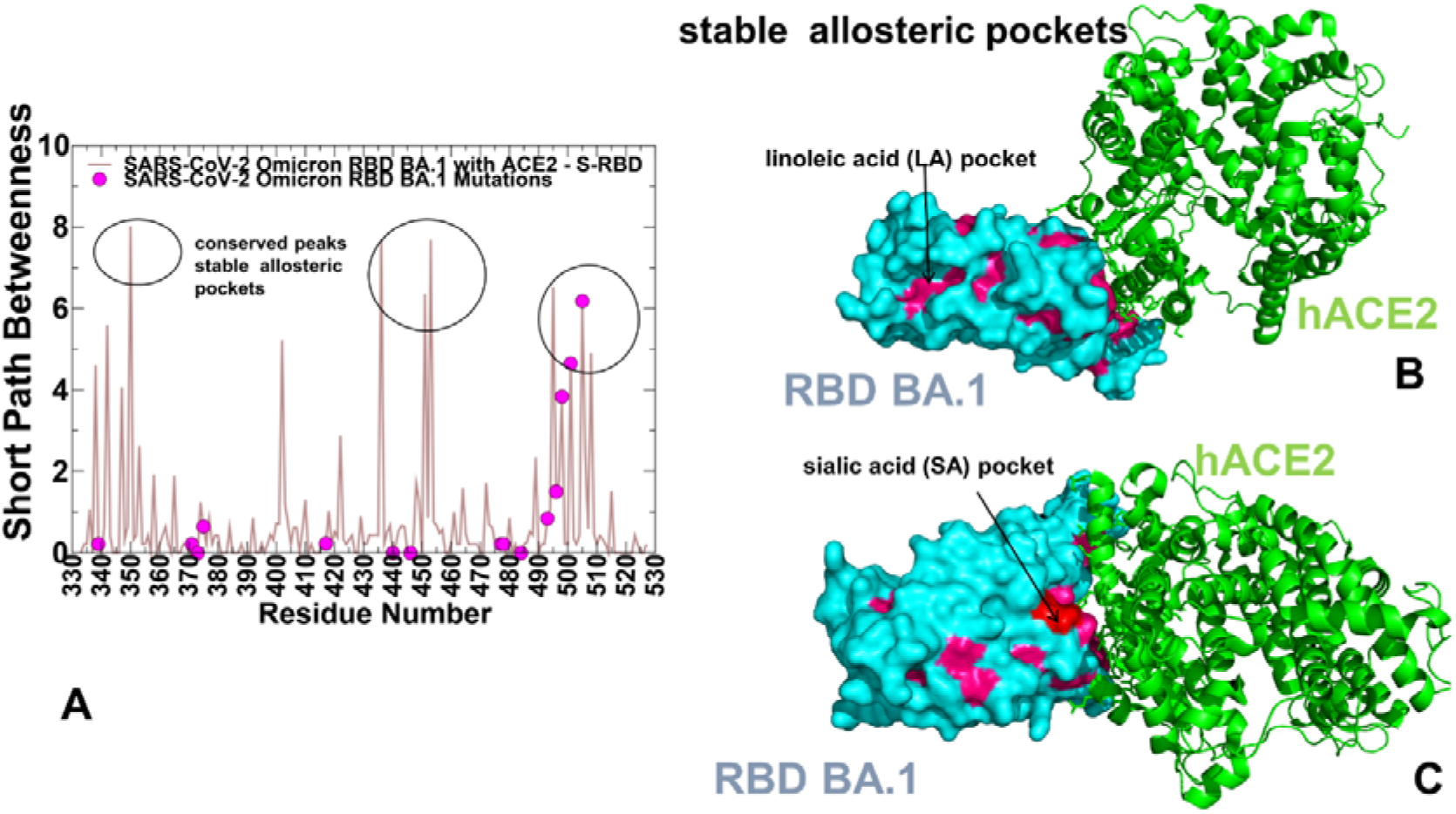
The dynamic network-based analysis of the Omicron RBD BA.1-hACE2 complex. (A) The residue-based short path betweenness (SPC) centrality profile. The centrality profiles of the S-RBD residues are shown in light brown lines and the positions of the BA.1 mutations are highlighted in magenta-colored filled circles. (B) Structural mapping of the SPC distribution peak clusters unveils potential allosteric binding pockets that are anchored by the allosteric hotspots. The RBD is shown in cyan surface and hACE2 is shown in green ribbons. The RBD pocket regions associated with network-mediating positions (high SPC values) are shown in pink surface. The SPC major peaks correspond the experimentally discovered allosteric binding site of linoleic acid [115]. This validated allosteric pocket is indicated by arrow and annotated. (C)) Structural mapping of the residues with high SPC values from regions different from the top clusters revealed the experimentally suggested allosteric site for sialic acid [116] (shown in red surface). Other more variable pockets are shown in pink surface.

We first examined the SPC distribution for the reference Omicron BA.1 complex with hACE2 (Figure 10) displaying several major cluster peaks associated with the residues involved in mediating global network connectivity and long-range communication in the system. The emergence of local clusters of mediating residues implies a high level of connectivity in the residue interaction network which may allow for diverse communication routes in the complex. Another interesting feature of the SPC distribution was the presence of several largest peaks that are associated with the RBD residues located in different regions (Figure 10A). More specifically, these single peaks corresponded to the F338, V341, F342, V350, I402 and W436 from the RBD core as well as Y451, Y453, Y495 and H505 (Figure 10A). Accordingly, these positions anchor clusters of allosteric centers that mediate long-range interactions between distal RBD regions and the RBM binding site. Importantly, the two most dominant allosteric clusters include residues in the RBD core that are also aligned with stability centers including V341, F342, V350, and W436 (Figure 10). The most striking finding of this network analysis is the fact that a major group of hydrophobic allosteric centers from RBD core (F338, V341 F342, W436) form a well-defined conserved allosteric binding pocket (Figure 10B) and this pocket corresponds precisely to the experimentally determined allosteric site where the essential free fatty acid linoleic acid (LA) binds [115].

The cryo-EM structure of the SARS-CoV-2 S linoleic acid complex showed a strong occupancy for this distal binding site and experimentally validated allosteric binding of LA to the S-RBD that stabilizes S conformation [115]. Furthermore, a similar pocket is shared in the structures of SARS-CoV and Middle East respiratory syndrome coronavirus (MERS-CoV) [115]. Based on the network-based reversed allosteric communication analysis, our results demonstrated that residues forming the conserved allosteric pocket contribute decisively to the inter-residue communications and form a major allosteric hotspot cluster (Figure 10A,B). Another cluster of network-mediating residues corresponded to Y451, Y453, Y495, Y489, Y501 and H505 residues that are involved in direct contacts with the hACE2 and also form the interfacial community cliques associated with potential epistatic effects. These findings implied that major allosteric communications occur between the conserved LA allosteric pocket and patches of the RBD interface anchored by Y453, Y489 and Y501 binding hotspots (Figure 10 A,B). By highlighting the Omicron mutational sites on the SPC distribution, we observed that most of the Omicron positions with the notable exception of R493, R498, Y501 and H505 are characterized by very small SPC values, suggesting that mutations and perturbations in these sites may not have a significant effect on long-range communications in the complex (Figure 10A). At the same time, the distribution highlighted a significant role of the RBD binding patch anchored by R493, R498 and Y501 in mediating allostery. Instructively, a complete mapping of the RBD residues displaying high SPC values demonstrated that these sites tend to occupy other well-defined pockets on the RBD surface that form “pocket-based” pathways connecting the LA allosteric site with the RBD binding interface patches (Figure 10B). One of these shallow pockets includes R03, D405 and R408 residues which form a portion of the inter-protomer allosteric site for N-Acetylneuraminic acid (Neu5Ac), a type of predominant sialic acid (Figure 10C) [116]. The enlarged allosteric site for this molecule includes S375 an K378 of other protomers [116]. In our study, we analyzed only RBD complexes, but the results suggested that the reversed allosteric mapping could also locate less prominent secondary pockets that may become more relevant in the context of the S trimer structure.

To provide a comprehensive allosteric analysis of the Omicron RBD complexes, we constructed the ensemble-based dynamic residue networks and computed the SPC distributions (Figure 11A-C). In this analysis, we examined similarities and differences between the SPC profiles and followed up with the reversed allosteric communication analysis to characterize allosteric binding pockets from mapping of the allosteric hotspots. In general, the distributions are similar, featuring peaks that corresponded to several conserved clusters of mediating centers shared among all Omicron variants : F338, V341, F342, V350, I402 and W436 the RBD core, Y451 and Y453, as well as binding site residues R493, Y495, R498, Y501 and H505 (Figure 11A-C). However, the intensity of these peaks were different among Omicron RBD complexes. Interestingly, we observed the largest cluster peaks in the RBD BA.2 complex (Figure 11B), suggesting that allosteric couplings between these regions may be stronger in the BA.2 complex. The distributions also highlighted a stronger peak for the cluster of the binding interface residues (R493, Y495, R498, Y501 and H505) for all Omicron subvariants (Figure 11A-C). The distribution peak around Y451 and Y453 is also pronounced for all Omicron RBD complexes, which is also related to the integrating role of these residue in mediating the interfacial communities and non-additive contributions.

**Figure 10.**
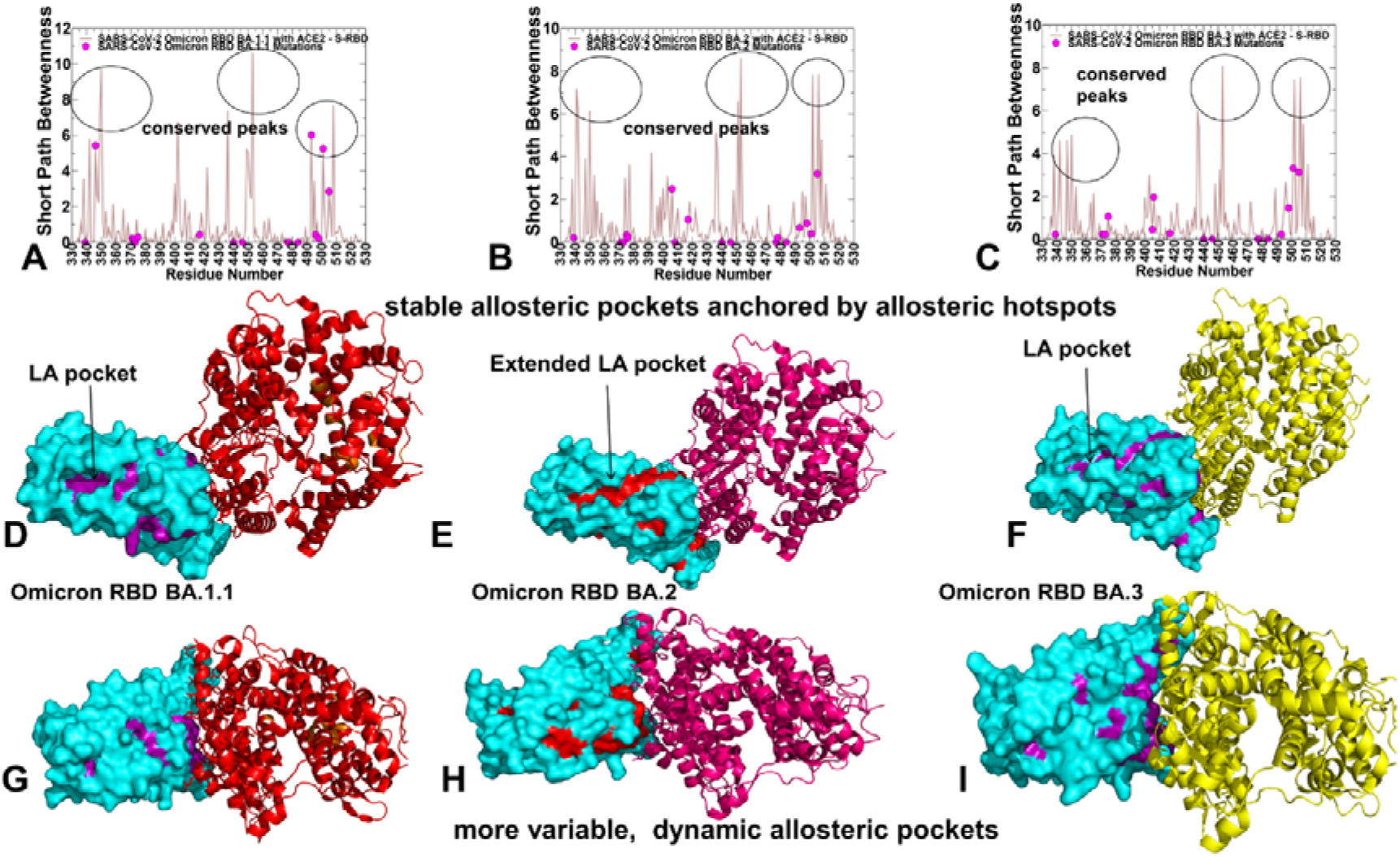
The dynamic network-based analysis of the Omicron RBD subvariant complexes with hACE2. The residue-based SPC centrality profiles of the S-RBD residues for the RBD BA.1.1-hACE2 complex (A), RBD BA.2-hACE2 complex (B) and RBD BA.3-hACE2 complex (C). The distributions are shown in light brown lines and the positions of the Omicron subvariant mutations are highlighted in magenta-colored filled circles. Structural mapping of the SPC distribution peak clusters unveils stable allosteric binding pockets that are anchored by the allosteric hotspots for the RBD BA.1.1-hACE2 complex. The RBD is shown in cyan surface, the pocket regions formed by mediating hotspots are shown in magenta surface and hACE2 is shown in red ribbons. The position of conserved LA allosteric pocket is indicated by an arrow (D). Structural mapping of the SPC distribution peak clusters for the RBD BA.2-hACE2 complex. The RBD is shown in cyan surface, the pocket regions formed are shown in red surface and hACE2 is shown in pink ribbons (E). Structural mapping of the SPC distribution peak clusters and distribution of allosteric pockets for the RBD BA.3-hACE2 complex. The RBD is shown in cyan surface, the pocket regions formed are shown in magenta surface and hACE2 is shown in green ribbons (F). (G-I) Structural mapping of the residues with high SPC values from regions different from the top clusters revealed more dynamic allosteric pockets. The annotations for the RBD, hACE2 and predicted pockets are same as in panels D-F.

By mapping RBD residues with high SPC values on the ensemble-average protein conformations, we observed consistent occupation of several primary conserved allosteric pockets (Figure 11D-F). Notably, these pockets are formed by residues from the three major cluster peaks. A common and most important characteristic of this mapping is that hydrophobic allosteric hotspots with the highest SPC values (F338, V341 F342, W436) are precisely aligned with the experimentally determined allosteric binding site [115]. Moreover, owing to some broadening of the SPC distribution for the BA.2 complex and greater number of high SPC positions, we detected that structural map of allosteric centers extended the LA pocket and illustrated a stronger inter-connectivity (Figure 11E). These results represent one of the important findings of the reversed allosteric communication analysis, showing that mediating centers of long-range interactions could anchor allosteric binding pockets and serve as proxy for identifying and ranking the importance of binding pockets for allosteric functions. Moreover, our analysis underscored “hidden” role of the binding energy hotspots R493, R498 and Y501 as potential mediators of epistatic effects and allosteric interactions in the Omicron RBD complexes. Remarkably, Omicron mutations in these key interface positions could fulfill multiple functional roles including the improved electrostatic interactions and enhanced complementarity with the host receptor, decisive contributions to the improved binding affinity with ACE2 by serving as binding energy hotspots, and additionally, acting as mediators of non-additive interfacial effects and allosteric communication hubs. We also detected multiple “secondary” binding pockets that are formed by residues with high SPC values but appreciably lower than the peaks of the distribution (Figure 11 G-I). In fact, these distributions are quite variable and specific for each of the Omicron variants. Indeed, the RBD residues that form these “minor” pockets are less significant in network terms and reflect a potential diversity in the ensembles of allosteric communication paths.

To summarize, by combining the dynamic network modeling with community-based assessment of potential epistatic effects and reversed allosteric analysis for mapping of allosteric pockets, we determined the indispensable role of R493, R498 and Y501 in mediating long-range communications beyond their established role as dominant binding hotspots. The results showed that using reversed allosteric mapping, we can identify the experimentally validated conserved allosteric binding pocket and characterize allosteric cross-talk between allosteric binding site and RBD-ACE2 binding interface. We found that major allosteric communications occur between the conserved LA allosteric pocket and patches of the RBD interface anchored by Y453, Y489 and Y501 binding hotspots.

## 3. Conclusions

In this study, we performed all-atom MD simulations of the RBD-ACE2 complexes for BA.1 BA.1.1, BA.2, an BA.3 Omicron subvariants, conducted a systematic mutational scanning of the RBD-ACE2 binding interfaces and analysis of electrostatic effects. The binding free energy computations of the Omicron RBD-ACE2 complexes and comprehensive examination of the electrostatic interactions quantify the driving forces of binding. A systematic mutational scanning of the RBD residues determined binding energy hotpots in the Omicron RBD-ACE2 complexes. By combining insights obtained from binding affinity analysis and mutational scanning, we showed that the enhanced binding affinity of the BA.2 complex may be determined by the strengthened inter-molecular electrostatic interactions mediated by R493, R498 and Y501 as well as cumulative contributions distributed over larger binding interface with the hACE2 receptor. Using the ensemble-based global network analysis, we proposed a community-based topological model of the Omicron RBD complexes to characterize the role of Omicron mutational sites in mediating non-additive epistatic effects of mutations. Our findings suggest that the extent of non-additive contributions to the binding affinity may be greater for the Omicron BA.1.1 and especially BA.2 complex that also featured the strongest binding affinity among the Omicron subvariants. We propose the network-centric adaptation of the reversed allosteric communication approach to identify allosteric mediating hotspots and characterize the distribution of allosteric binding pockets on the RBD. These results showed that mediating centers of long-range interactions could anchor allosteric binding pockets and serve as proxy for identifying and ranking the importance of binding pockets for allosteric functions. Moreover, our analysis underscored “hidden” role of the binding energy hotspots R493, R498 and Y501 as important mediators of epistatic effects and allosteric interactions in the Omicron RBD complexes. Remarkably, Omicron mutations in these key interface positions could fulfill multiple functional roles including the improved electrostatic interactions and enhanced complementarity with the host receptor, decisive contributions to the improved binding affinity with ACE2 by serving as binding energy hotspots, and additionally, acting as mediators of non-additive interfacial effects and allosteric communication hubs.

## 4. Materials and Methods

### 4.1. Molecular Dynamics Simulations

All structures were obtained from the Protein Data Bank [117]. During structure preparation stage, protein residues in the crystal structures were inspected for missing residues and protons. Hydrogen atoms and missing residues were initially added and assigned according to the WHATIF program web interface [118]. The missing loops in the studied cryo-EM structures of the SARS-CoV-2 S protein were reconstructed and optimized using template-based loop prediction approaches ModLoop [119] and ArchPRED server [120]. The side chain rotamers were refined and optimized by SCWRL4 tool [121]. The protein structures were then optimized using atomic-level energy minimization with a composite physics and knowledge-based force fields as implemented in the 3Drefine method [122]. The atomistic structures from simulation trajectories were further elaborated by adding N-acetyl glycosamine (NAG) glycan residues and optimized. Using NAMD 2.13-multicore-CUDA package [123] with CHARMM36 force field [124], 10 independent all-atom MD simulations (500 ns each simulation) were carried out for each of the Omicron RBD-hACC2 complexes : RBD BA.1-hACE2 (pdb id 7WBP), S-RBD RBD BA.1.1-hACE2 (pd id 7XAZ), RBD BA.2-hACE2 (pdb id 7XB0), and RBD BA.3-hACE2 (pdb id 7XB1) (Table 2). The equilibrium ensembles for the analysis are derived by aggregating the 10 independent MD trajectories for every system. The structures of the SARS-CoV-2 S-RBD complexes were prepared in Visual Molecular Dynamics (VMD 1.9.3) [125] by placing them in a TIP3P water box with 20 Å thickness from the protein. Assuming normal charge states of ionizable groups corresponding to pH = 7, sodium (Na+) and chloride (Cl-) counter-ions were added to achieve charge neutrality and a salt concentration of 0.15 M NaCl was maintained. All Na^+^ and Cl^-^ ions were placed at least 8 Å away from any protein atoms and from each other. The long-range non-bonded van der Waals interactions were computed using an atom-based cutoff of 12 Å with the switching function beginning at 10 Å and reaching zero at 14 Å. SHAKE method was used to constrain all bonds associated with hydrogen atoms. Simulations were run using a leap-frog integrator with a 2 fs integration time step. ShakeH algorithm of NAMD was applied for water molecule constraints. The long-range electrostatic interactions were calculated using the particle mesh Ewald method [126] with a cut-off of 1.0 nm and a fourth order (cubic) interpolation. Simulations were performed under NPT ensemble with Langevin thermostat and Nosé-Hoover Langevin piston at 310 K and 1 atm. The damping coefficient (gamma) of the Langevin thermostat was 1/ps. The Langevin piston Nosé-Hoover method in NAMD is a combination of the Nose-Hoover constant pressure method [127] with piston fluctuation control implemented using Langevin dynamics [128,129]. Energy minimization was conducted using the steepest descent method for 100,000 steps. All atoms of the complex were first restrained at their crystal structure positions with a force constant of 10 Kcal mol^-1^ Å^-2^. Equilibration was done in steps by gradually increasing the system temperature in steps of 20K starting from 10K until 310 K and at each step 1ns equilibration was done keeping a restraint of 10 Kcal mol-1 Å-2 on the protein C_α_ atoms. After the restrains on the protein atoms were removed, the system was equilibrated for additional 10 ns. An NPT production simulation was run on the equilibrated structures for 500 ns keeping the temperature at 310 K and constant pressure (1 atm).

### 4.2. Electrostatic Calculations

In the framework of continuum electrostatics, the electrostatic potential *φ* for biological macromolecules can be obtained by solving the Poisson–Boltzmann equation (PBE)

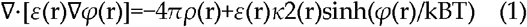

where *φ* (r) is the electrostatic potential, *ε*(r) is the dielectric distribution, *ρ*(r) is the charge density based on the atomic structures, *κ* is the Debye–Huckel parameter, kB is the Boltzmann constant, and T is the temperature. The electrostatic interaction potentials are computed for the averaged RBD-hACE2 conformations using the APBS-PDB2PQR software [84,85] based on the Adaptive Poisson–Boltzmann Solver (APBS) [84] and visualized using the VMD visualization tool [86]. These resources are available from the APBS/PDB2PQR website: http://www.poissonboltzmann.org/. The atomic charges and radii are assigned in this approach based on the chosen force field. For consistency with MD simulations, we opted in PDB2PQR for parameters from CHARMM22 [130]. Using DelPhiPKa protonation and pKa calculations [88,89], we also computed the electrostatic polar energies and desolvation energies for the RBD residues based on the equilibrium structures of the Omicron RBD-hACE2 complexes.

### 4.3. Binding Free Energy Computations

The binding free energies were computed for the Omicron RBD-hACE2 complexes using the MM-PBSA method [81] using the pipeline tool Calculation of Free Energy (CaFE) implemented as VMD plugin [82]. For these calculations, 1,000 simulation samples from MD trajectories were used for each of the studied complexes. The entropy variations in the binding free energy computations were not considered due to high structural and dynamic similarities between Omicron RBD-hACE2 complexes. This approach is consistent with previous MM-PBSA computations of binding affinity for the RBD-hACE2 complexes [59]. The binding free energy computations were also performed using a contact-based predictor of binding affinity Prodigy [83], where computations of the binding affinity scores were averaged over 1,000 simulation samples from MD trajectories.

### 4.4. Mutational Scanning and Sensitivity Analysis

We conducted mutational scanning analysis of the binding epitope residues for the SARS-CoV-2 S protein complexes. Each binding epitope residue was systematically mutated using all substitutions and corresponding protein stability changes were computed. BeAtMuSiC approach [93-95] was employed that is based on statistical potentials describing the pairwise inter-residue distances, backbone torsion angles and solvent accessibilities, and considers the effect of the mutation on the strength of the interactions at the interface and on the overall stability of the complex. The binding free energy of protein-protein complex can be expressed as the difference in the folding free energy of the complex and folding free energies of the two protein binding partners:

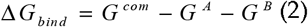

The change of the binding energy due to a mutation was calculated then as the following

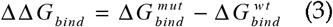

We leveraged rapid calculations based on statistical potentials to compute the ensemble-averaged binding free energy changes using equilibrium samples from simulation trajectories. The binding free energy changes were computed by averaging the results over 1,000 equilibrium samples for each of the studied systems.

### 4.5. Dynamic Network Analysis and Topological Clique-Based Model for Assessment of Non-Additivity

A graph-based representation of protein structures [131,132] is used to represent residues as network nodes and the inter-residue edges to describe non-covalent residue interactions. The network edges that define residue connectivity are based on non-covalent interactions between residue side-chains. The residue interaction networks were constructed by incorporating the topology-based residue connectivity MD-generated maps of residues cross-correlations [102] and coevolutionary couplings between residues measured by the mutual information scores [103]. The edge lengths in the network are obtained using the generalized correlation coefficients ***R***_*Ml*_(***X***_***i***_, ***X***_***J***_)associated with the dynamic correlation and mutual information shared by each pair of residues. The length (i.e. weight) *w*_*ij*_ = − log[***R***_*Ml*_(***X***_***i***_, ***X***_***J***_)] of the edge that connects nodes ***i*** and ***j*** is defined as the element of a matrix measuring the generalized correlation coefficient ***R***_*Ml*_(***X***_***i***_, ***X***_***J***_) as between residue fluctuations in structural and coevolutionary dimensions. Network edges were weighted for residue pairs with ***R***_*Ml*_(***X***_***i***_, ***X***_***J***_) > 0.5 in at least one independent simulation as was proposed in our earlier study [76]. The matrix of communication distances is obtained using generalized correlation between composite variables describing both dynamic positions of residues and coevolutionary mutual information between residues. As a result, the weighted graph model defines a residue interaction network that favors a global flow of information through edges between residues associated with dynamics correlations and coevolutionary dependencies.

The Residue Interaction Network Generator (RING) program was employed for the initial generation of residue interaction networks based on the single structure [133] and the conformational ensemble [134] where edges have an associated weight reflecting the frequency in which the interaction present in the conformational ensemble. The residue interaction network files in xml format were obtained for all structures using RING v3.0 webserver freely available at https://ring.biocomputingup.it/submit. Network graph calculations were performed using the python package NetworkX [135]. Using the constructed protein structure networks, we computed the residue-based betweenness parameter. The short path betweenness of residue *i* is defined to be the sum of the fraction of shortest paths between all pairs of residues that pass through residue *i:*

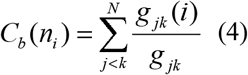

where *g*_*jk*_ denotes the number of shortest geodesics paths connecting *j* and k, and *g*_*jk*_ (*i*) is the number of shortest paths between residues *j* and k passing through the node *n*_*i*_. Residues with high occurrence in the shortest paths connecting all residue pairs have a higher betweenness values. For each node *n*, the betweenness value is normalized by the number of node pairs excluding *n* given as (*N* - 1)(*N* - 2) / 2, where *N* is the total number of nodes in the connected component that node *n* belongs to. The normalized short path betweenness of residue *i* can be expressed as follows:

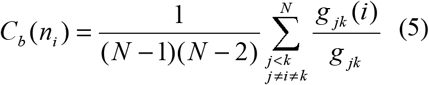

*g*_*jk*_ is the number of shortest paths between residues *j* and k; *g*_*jk*_ (*i*) is the fraction of these shortest paths that pass through residue *i*.

The Girvan-Newman algorithm [136,137] is used to identify local communities. In this approach, edge centrality (also termed as edge betweenness) is defined as the ratio of all the shortest paths passing through a particular edge to the total number of shortest paths in the network. The method employs an iterative elimination of edges with the highest number of the shortest paths that go through them. By eliminating edges, the network breaks down into smaller communities. The analysis of the interaction networks was done using network parameters such as cliques and communities. The *k*-cliques are complete sub graphs of size *k* in which each node is connected to every other node. In our application, a *k*-clique is defined as a set of *k* nodes that are represented by the protein residues in which each node is connected to all the other nodes. A *k*-clique community is determined by the Clique Percolation Method [138] as a subgraph containing *k*-cliques that can be reached from each other through a series of adjacent k-cliques. We have used a community definition according to which in a *k*-clique community two *k*-cliques share *k*−1 or *k*−2 nodes. Computation of the network parameters was performed using the Clique Percolation Method as implemented in the CFinder program [139]. Given the chosen interaction cutoff ***I***_min_ we typically obtain communities formed as a union of *k* =3 and *k* =4 cliques. The interaction cliques and communities were considered to be dynamically stable if these interaction networks remained to be intact in more than 75% of the ensemble conformations.

The stable 3-cliques that are formed by the interfacial RBD and hACE2 residues are determined for each of the studied Omicron RBD-hACE2 complexes. We then compute *P*_*ab*_ probabilities for all sites (*a, b*) to be held together in a 3-clique due to either direct or indirect interactions. The interfacial sites that belong to the same clique and display high *P*_*ab*_ probability are assumed to have propensity for stronger non-additive effects.

