## Supplementary Tables S1-S6 for "Atomistic Simulations and Network-Based Energetic Profiling of Binding and Allostery in the SARS-CoV-2 Spike Omicron BA.1, BA.1.1, BA.2 and BA.3 Subvariant Complexes with the Host Receptor: Revealing Hidden Functional Roles of the Binding Hotspots in Mediating Epistatic Effects and Long-Range Commun"

Received: date; Accepted: date; Published: date

**Table S1.** Statistical analysis of the intermolecular contact residues in Omicron RBD-hACE2 complexes.*

| hACE2 | BA.1 RBD | BA.1.1 RBD | BA.2 RBD | BA.3 RBD |
|---|---|---|---|---|
| S19 | A475, G476,N477 | A475, G476,N477 | A475, G476,N477 | A475, G476,N477 |
| T20 | | N477 | N477 | A475,N477 |
| Q24 | A475, G476,N477 F486, N487, Y489 | A475, G476,N477 F486, N487, Y489 | A475, G476,N477 F486,N487, Y489 | A475, G476,N477 F486,N487, Y489 |
| T27 | F456, Y473, A475,Y489 | F456, Y473, A475,Y489 | F456, Y473, A475,Y489 | F456, Y473, A475,Y489 |
| F28 | N487,Y489 | N487,Y489 | N487,Y489 | Y489 |
| F30 | L455, F456 | N417, L455, F456 | N417, L455, F456 | L455, F456 |
| K31 | L455, F456, Y489,R493 | L455, F456, Y489,F490, L492, R493 | L455, F456, G485,Y489, R493 | L455, F456, Y489,R493 |
| H34 | Y453, L455, R493, S494, Y495 | N417, Y453, L455, R493 | R403, N417, Y453, L455, R493 | N417, Y453, L455, R493 |
| E35 | R493 | R493 | R493 | R493 |
| E37 | H505 | H505 | H505 | H505 |
| D38 | Y449, S496, R498, Y501 | Y449, R493, S494, Y495, S496, R498, Y501 | Y449, Y495, G496, R498, Y501 | Y449, Y495, R498, Y501 |
| Y41 | R498, T500, Y501 | R498, T500, Y501 | R498, T500, Y501 | R498, T500, Y501 |
| Q42 | S446, Y449, R498 | Y449, R498 | Y449, R498 | Y449, R498 |
| L45 | R498,T500 | V445, R498,T500 | V445, R498,T500 | V445, R498,T500 |
| L79 | F486 | F486 | G485,F486 | F486 |
| M82 | F486 | F486 | F486 | F486 |
| Y83 | F486, N487, Y489 | F486, N487, Y489 | F486, N487, Y489 | F486, N487, Y489 |
| Q325 | V593 | | Q506 | V503, Q506 |

| | | | | |
|---|---|---|---|---|
| G326 | | | | T500 |
| N330 | T500 | T500 | P499,T500 | P499,T500 |
| G352 | | | Y501,G502 | Y501,G502 |
| K353 | R403, Y495, S496, T500, Y501, G502, H505 | Y495,S496 T500, Y501, G502,H505 | R403, Y495, T500,Y501, G502, V503, H505 | R403, Y495, T500,Y501, G502, V503, H505 |
| G354 | T500,Y501, G502,V503, H505 | T500, Y501, G502, H505 | T500,Y501, G502,V503, H505 | T500,Y501, G502,V503, H505 |
| D355 | T500, Y501,G502 | T500, Y501,G502 | T500, Y501,G502 | T500, Y501,G502 |
| R357 | | T500 | T500 | |

*Two residues are defined in contact if any of their heavy atom is within a distance of 5.0 Å

**Table S2.** The Occupancy of the Pairwise Interactions in the Omicron RBD-hACE2 Complexes

| | Interaction | BA.1-ACE2 | BA.1.1-ACE2 | BA.2-ACE2 | BA.3-ACE2 |
|---|---|---|---|---|---|
| Salt bridges | R403-E37 | 65% | 78% | 73% | 73% |
| | K440-E329 | 31% | 56% | 54% | 54% |
| | R493-E35 | 77% | 88% | 92% | 99% |
| | R493-D38 | 26% | 95% | 89% | 89% |
| | R498-D38 | 59% | 97% | 95% | 83% |
| Hydrophobic Interactions | F456-T27 | 95% | 87% | 96% | 88% |
| | Y473-T27 | 92% | 88% | 89% | 85% |
| | A475-T27 | 88% | 87% | 93% | 83% |
| | F486-F28 | 78% | 92% | 97% | 90% |
| | F486-L79 | 85% | 93% | 89% | 82% |
| | F486-M82 | 85% | 92% | 96% | 90% |
| | F486-Y83 | 90% | 98% | 95% | 87% |
| | Y489-F28 | 97% | 90% | 94% | 95% |
| | Y489-L79 | 90% | 89% | 95% | 86% |
| | Y489-Y83 | 96% | 78% | 82% | 88% |
| Hydrogen Bonds | Y453-H34 | 36% | 42% | 32% | 92% |
| | Y449-D38 | 45% | 54% | 52% | 58% |
| | A475-S19 | 30% | 70% | 65% | 85% |
| | N477-S19 | 28% | 72% | 77% | 97% |
| | N487-Y83 | 42% | 76% | 82% | 92% |
| | T500-D355 | 62% | 74% | 77% | 90% |
| | T500-Y41 | 62% | 82% | 80% | 95% |
| | G502-K353 | 78% | 82% | 84% | 78% |
| | Y501-K353 | 66% | 92% | 90% | 84% |
| Specific Interactions | Y501-Y41 | 85 | 88% | 92% | 91% |
| | Y501-K353 | 82% | 88% | 96% | 92% |

**Table S3.** The electrostatic polar energies and desolvation energies for the RBD residues based on the equilibrium structure of the Omicron RBD BA.1-hACE2 complex.*

| Residue Name | pKa | Elec.polar G(charged) | Elec. polar G(neutral) | Desolvation G(charged) | Desolvation G(neutral) |
|---|---|---|---|---|---|
| ARG0346 | 12.19 | -0.1359 | 0.0049 | 0.0361 | 0.0050 |
| ARG0355 | 12.97 | -0.1693 | 0.0852 | 0.0967 | 0.0211 |
| LYS0356 | 11.01 | -0.1282 | 0.0403 | 0.0381 | 0.0060 |
| ARG0357 | 12.26 | -0.0860 | 0.0602 | 0.0467 | 0.0081 |
| ASP0364 | 3.80 | -0.2422 | -0.0248 | 0.0870 | 0.0284 |
| LYS0378 | 10.80 | -0.0449 | 0.0139 | 0.0162 | 0.0076 |
| LYS0386 | 10.87 | -0.0564 | -0.0173 | 0.0086 | 0.0044 |
| ASP0389 | 3.92 | 0.0681 | 0.0532 | 0.0841 | 0.0318 |
| ASP0398 | 2.74 | 0.1148 | 0.0403 | 0.2053 | 0.0562 |
| ARG0403 | 13.09 | 0.2182 | 0.0144 | 0.1298 | 0.0228 |
| ASP0405 | 2.85 | -0.2475 | 0.0219 | 0.0930 | 0.0287 |
| GLU0406 | 2.41 | -0.5120 | -0.1416 | 0.1728 | 0.0814 |
| ARG0408 | 12.74 | -0.1853 | 0.1030 | 0.0618 | 0.0084 |
| ASP0420 | 3.16 | -0.2342 | -0.0468 | 0.1100 | 0.0307 |
| LYS0424 | 11.22 | -0.2218 | -0.0781 | 0.0839 | 0.0091 |
| ASP0427 | 3.49 | -0.1535 | 0.0115 | 0.0506 | 0.0206 |
| ASP0428 | 3.59 | -0.3138 | -0.0418 | 0.0885 | 0.0312 |
| LYS0440 | 10.79 | 0.0707 | 0.0037 | 0.0147 | 0.0045 |
| ASP0442 | 2.31 | -0.1857 | 0.0955 | 0.1677 | 0.0415 |
| LYS0444 | 10.48 | 0.0411 | -0.0111 | 0.0447 | 0.0045 |
| ARG0454 | 12.94 | -0.8003 | -0.1488 | 0.2183 | 0.0326 |
| ARG0457 | 13.06 | -0.2267 | -0.0202 | 0.1139 | 0.0284 |
| LYS0458 | 11.08 | -0.1643 | 0.0370 | 0.0328 | 0.0053 |
| LYS0462 | 10.93 | -0.0337 | 0.0037 | -0.0013 | 0.0051 |
| GLU0465 | 3.06 | -0.1407 | 0.0048 | 0.0779 | 0.0053 |
| ARG0466 | 12.46 | -0.5530 | -0.2125 | 0.0913 | 0.0274 |
| ASP0467 | 2.77 | -0.3815 | -0.0474 | 0.1338 | 0.0413 |
| GLU0471 | 3.55 | 0.0021 | 0.0162 | 0.0232 | 0.0014 |
| LYS0478 | 10.82 | -0.0092 | 0.0056 | 0.0107 | 0.0046 |
| ARG0493 | 12.97 | -0.2343 | 0.0692 | 0.1424 | 0.0254 |
| ARG0498 | 13.24 | -0.3585 | 0.0178 | 0.1809 | 0.0226 |
| HIS0505 | 5.98 | 0.1497 | -0.0646 | 0.0853 | 0.0138 |
| ARG0509 | 12.97 | -0.3253 | 0.1119 | 0.1667 | 0.0291 |
| GLU0516 | 3.79 | 0.0190 | -0.0273 | 0.0752 | 0.0050 |
| HIS0519 | 6.52 | -0.0064 | 0.0024 | 0.0128 | 0.0039 |

* pKa value for each ionizable residue with associated energy terms (the unit here is kcal/mol). The electrostatic polar energy for individual residue in its protonated (+/-) state and neutral state, and the desolvation energy for individual residue in its protonated (+/-) state and neutral state.

**Table S4.** The electrostatic polar energies and desolvation energies for the RBD residues based on the equilibrium structure of the Omicron RBD BA.1.1-hACE2 complex.*

| Residue Name | pKa | Elec. polar G(charged) | Elec. polar G(neutral) | Desolvation G(charged) | Desolvation G(neutral) |
|---|---|---|---|---|---|
| ASP0339 | 3.81 | -0.0908 | 0.0711 | 0.0541 | 0.0211 |
| GLU0340 | 3.46 | -0.0170 | 0.0221 | 0.0496 | 0.0037 |
| LYS0346 | 10.77 | -0.0487 | -0.0048 | 0.0129 | 0.0039 |
| ARG0355 | 13.03 | -0.2704 | 0.0674 | 0.1592 | 0.0396 |
| LYS0356 | 11.07 | -0.1082 | 0.0108 | 0.0445 | 0.0048 |
| ARG0357 | 12.15 | -0.0903 | -0.0354 | 0.0661 | 0.0146 |
| ASP0364 | 3.50 | -0.5453 | -0.2062 | 0.1596 | 0.3294 |
| LYS0378 | 10.79 | -0.0481 | 0.0023 | 0.0283 | 0.0063 |
| LYS0386 | 10.76 | 0.2961 | 0.0784 | 0.0457 | 0.0072 |
| ASP0389 | 3.97 | 0.2925 | 0.0105 | 0.0896 | 0.0277 |
| ASP0398 | 2.69 | 0.2209 | 0.0373 | 0.2015 | 0.0707 |
| ARG0403 | 13.00 | 0.2234 | 0.0133 | 0.1117 | 0.0170 |
| ASP0405 | 2.79 | -0.2302 | 0.0127 | 0.0779 | 0.0254 |
| GLU0406 | 2.55 | -0.2820 | 0.0030 | 0.1709 | 0.0287 |
| ARG0408 | 12.56 | -0.0134 | 0.0027 | 0.0388 | 0.0084 |
| ASP0420 | 3.19 | -0.2075 | 0.0623 | 0.1201 | 0.0455 |
| LYS0424 | 11.52 | -0.2984 | 0.0286 | 0.0807 | 0.0075 |
| ASP0427 | 3.52 | -0.1363 | 0.0068 | 0.0576 | 0.0222 |
| ASP0428 | 3.79 | -0.2309 | -0.0258 | 0.0887 | 0.0318 |
| LYS0440 | 10.79 | 0.0684 | 0.0046 | 0.0047 | 0.0050 |
| ASP0442 | 2.50 | -0.2114 | 0.1357 | 0.1722 | 0.0451 |
| LYS0444 | 10.79 | -0.1484 | -0.0160 | 0.0476 | 0.0039 |
| ARG0454 | 12.89 | -0.5339 | 0.0827 | 0.1961 | 0.0331 |
| ARG0457 | 13.07 | -0.3141 | -0.0174 | 0.1126 | 0.0206 |
| LYS0458 | 11.04 | -0.0493 | 0.0117 | 0.0201 | 0.0045 |
| LYS0462 | 11.00 | -0.0455 | 0.0027 | 0.0230 | 0.0053 |
| GLU0465 | 2.95 | -0.1673 | -0.0385 | 0.0817 | 0.0064 |
| ARG0466 | 12.49 | -0.3143 | 0.0291 | 0.0609 | 0.0109 |
| ASP0467 | 2.81 | -0.3548 | -0.0445 | 0.1265 | 0.0396 |
| GLU0471 | 3.50 | -0.0063 | -0.0032 | 0.0246 | 0.0020 |
| LYS0478 | 10.75 | 0.0994 | -0.0091 | 0.0164 | 0.0053 |
| ARG0493 | 13.62 | -0.2106 | -0.0522 | 0.1110 | 0.0240 |
| ARG0498 | 12.55 | -0.3407 | -0.0812 | 0.1123 | 0.0127 |
| HIS0505 | 6.02 | 0.1086 | -0.0701 | 0.0749 | 0.0119 |
| ARG0509 | 13.06 | -0.4417 | 0.1141 | 0.1452 | 0.0320 |
| GLU0516 | 3.50 | -0.1058 | -0.0044 | 0.0568 | 0.0027 |
| HIS0519 | 6.71 | 0.1577 | 0.0439 | 0.0428 | 0.0138 |

**Table S5.** The electrostatic polar energies and desolvation energies for the RBD residues based on the equilibrium structure of the Omicron RBD BA.2-hACE2 complex.*

| Residue Name | pKa | Elec.polar G(charged) | Elec. polar G(neutral) | Desolvation G(charged) | Desolvation G(neutral) |
|---|---|---|---|---|---|
| ASP0339 | 3.82 | -0.0993 | 0.0421 | 0.0832 | 0.0277 |
| GLU0340 | 3.05 | -0.0686 | 0.0293 | 0.0811 | 0.0220 |
| ARG0346 | 12.14 | -0.0972 | -0.0166 | 0.0460 | 0.0122 |
| ARG0355 | 13.06 | -0.1927 | 0.0874 | 0.1973 | 0.0739 |
| LYS0356 | 11.20 | 0.0111 | -0.0106 | 0.1754 | 0.1047 |
| ARG0357 | 12.25 | -0.3185 | -0.1527 | 0.1040 | 0.0301 |
| ASP0364 | 3.52 | -0.5235 | -0.0363 | 0.1045 | 0.0360 |
| LYS0378 | 10.95 | -0.2034 | 0.0732 | 0.0501 | 0.0055 |
| LYS0386 | 10.87 | 0.0003 | -0.0047 | 0.0097 | 0.0051 |
| ASP0389 | 3.91 | 0.0406 | 0.0214 | 0.0672 | 0.0237 |
| ASP0398 | 2.20 | 0.1758 | 0.0302 | 0.2426 | 0.1627 |
| ARG0403 | 12.97 | 0.1549 | 0.0596 | 0.0951 | 0.0115 |
| GLU0406 | 2.51 | -0.3939 | -0.1179 | 0.1513 | 0.0139 |
| ASP0420 | 3.14 | -0.3525 | 0.0029 | 0.1144 | 0.0308 |
| LYS0424 | 11.82 | -0.4666 | 0.0532 | 0.1300 | 0.0151 |
| ASP0427 | 3.46 | -0.1642 | 0.0409 | 0.0713 | 0.0250 |
| ASP0428 | 3.61 | -0.3245 | -0.0656 | 0.0823 | 0.0297 |
| LYS0440 | 10.83 | -0.0327 | -0.0003 | 0.0110 | 0.0049 |
| ASP0442 | 2.21 | -0.3310 | 0.1723 | 0.2134 | 0.0540 |
| LYS0444 | 10.53 | -0.0529 | 0.0196 | 0.0383 | 0.0046 |
| ARG0454 | 13.09 | -0.9663 | -0.1073 | 0.2756 | 0.0419 |
| ARG0457 | 13.22 | -0.3610 | 0.0498 | 0.1616 | 0.0614 |
| LYS0458 | 11.01 | -0.0046 | 0.0223 | 0.0247 | 0.0053 |
| LYS0462 | 10.99 | -0.0913 | -0.0080 | 0.0292 | 0.0077 |
| GLU0465 | 2.79 | -0.2246 | -0.0936 | 0.1136 | 0.0122 |
| ARG0466 | 12.50 | -0.6164 | -0.1977 | 0.0791 | 0.0173 |
| ASP0467 | 2.92 | -0.2084 | -0.1494 | 0.1630 | 0.0588 |
| GLU0471 | 3.46 | -0.0586 | 0.0318 | 0.0229 | 0.0008 |
| LYS0478 | 10.82 | 0.0146 | 0.0120 | 0.0075 | 0.0058 |
| ARG0493 | 13.04 | -0.3137 | -0.1730 | 0.2313 | 0.0685 |
| ARG0498 | 13.46 | -0.6965 | 0.0474 | 0.2437 | 0.0374 |
| HIS0505 | 5.78 | 0.0517 | -0.0336 | 0.1003 | 0.0180 |
| ARG0509 | 13.04 | -0.4000 | 0.2088 | 0.2133 | 0.0454 |
| GLU0516 | 3.26 | -0.4601 | -0.0451 | 0.0956 | 0.0168 |
| HIS0519 | 6.50 | 0.0315 | -0.0072 | 0.0120 | 0.0046 |

**Table S6.** The electrostatic polar energies and desolvation energies for the RBD residues based on the equilibrium structure of the Omicron RBD BA.3-hACE2 complex.*

| Residue Name | pKa | Elec.polar G(charged) | Elec. polar G(neutral) | Desolvation G(charged) | Desolvation G(neutral) |
|---|---|---|---|---|---|
| ASP0339 | 3.78 | -0.1619 | 0.0544 | 0.0680 | 0.0228 |
| GLU0340 | 3.03 | -0.0815 | 0.0242 | 0.0640 | 0.0342 |
| ARG0346 | 12.15 | -0.1075 | -0.0152 | 0.0566 | 0.0178 |
| ARG0355 | 13.04 | -0.2072 | 0.0870 | 0.1497 | 0.0485 |
| LYS0356 | 11.12 | 0.0344 | 0.0037 | 0.0519 | 0.0172 |
| ARG0357 | 12.17 | -0.1901 | -0.2289 | 0.0486 | 0.0103 |
| ASP0364 | 3.91 | -0.0739 | 0.0332 | 0.1188 | 0.0345 |
| LYS0378 | 10.85 | -0.1407 | 0.0142 | 0.0316 | 0.0042 |
| LYS0386 | 10.91 | 0.0002 | -0.0080 | 0.0018 | 0.0043 |
| ASP0389 | 3.83 | 0.0469 | 0.0261 | 0.1047 | 0.0339 |
| ASP0398 | 2.50 | 0.1899 | 0.0363 | 0.2494 | 0.0886 |
| ARG0403 | 13.13 | 0.0608 | 0.1371 | 0.0939 | 0.0155 |
| GLU0406 | 2.22 | -0.5186 | -0.1393 | 0.1896 | 0.1051 |
| ARG0408 | 12.21 | -0.0873 | -0.0165 | 0.0176 | 0.0039 |
| ASP0420 | 3.32 | -0.1717 | 0.0651 | 0.1037 | 0.0263 |
| LYS0424 | 11.39 | -0.2473 | 0.0059 | 0.1422 | 0.0105 |
| ASP0427 | 3.50 | -0.1628 | 0.0256 | 0.0613 | 0.0219 |
| ASP0428 | 3.58 | -0.3356 | -0.0390 | 0.0924 | 0.0302 |
| LYS0440 | 10.85 | -0.1198 | 0.0033 | 0.0164 | 0.0037 |
| ASP0442 | 2.10 | -0.2708 | 0.0907 | 0.2122 | 0.0692 |
| LYS0444 | 10.45 | 0.0291 | -0.0179 | 0.0588 | 0.0063 |
| ARG0454 | 12.99 | -0.9878 | -0.1701 | 0.3992 | 0.0710 |
| ARG0457 | 13.20 | -0.4717 | -0.2062 | 0.1838 | 0.0345 |
| LYS0458 | 11.02 | 0.0111 | -0.0061 | 0.0520 | 0.0071 |
| LYS0462 | 11.02 | -0.0454 | -0.0015 | 0.0271 | 0.0050 |
| GLU0465 | 2.83 | -0.2698 | -0.0884 | 0.1026 | 0.0095 |
| ARG0466 | 12.52 | -0.6312 | -0.1788 | 0.0898 | 0.0226 |
| ASP0467 | 3.09 | 0.0036 | -0.2796 | 0.1853 | 0.1827 |
| GLU0471 | 3.42 | -0.0223 | 0.0278 | 0.0305 | 0.0017 |
| LYS0478 | 10.85 | -0.0366 | 0.0047 | 0.0026 | 0.0057 |
| ARG0493 | 13.00 | -0.1220 | 0.0565 | 0.1390 | 0.0270 |
| ARG0498 | 13.35 | -0.6293 | 0.0144 | 0.2198 | 0.0332 |
| HIS0505 | 5.76 | 0.0638 | 0.0190 | 0.0869 | 0.0140 |
| ARG0509 | 12.99 | -0.3698 | 0.1129 | 0.3041 | 0.0622 |
| GLU0516 | 3.50 | -0.2544 | 0.0114 | 0.1275 | 0.0238 |
| HIS0519 | 6.52 | -0.0329 | 0.0030 | 0.0136 | 0.0042 |
